# Abscisic acid and GIGANTEA signalling converge to regulate the recruitment of CONSTANS to the *FT* promoter and activate floral transition

**DOI:** 10.1101/2024.05.29.595696

**Authors:** Alice Robustelli Test, Giorgio Perrella, Beatrice Landoni, Sara Colanero, Aldo Sutti, Paolo Korwin Krukowski, Tianyuan Xu, Elisa Vellutini, Giulia Castorina, Massimo Galbiati, Damiano Martignago, Eirini Kaiserli, Chiara Tonelli, Lucio Conti

**Affiliations:** Dipartimento di Bioscienze, Università degli Studi di Milano, Italy; School of Molecular Biosciences, College, of Medical, Veterinary and Life Sciences, University of Glasgow, UK; Dipartimento di Scienze agrarie e ambientali, Università degli Studi di Milano, Italy; Istituto di Biologia e Biotecnologia Agraria – IBBA, CNR – Milano, Italy

**Keywords:** flowering time regulation, environmental and stress responses, florigen signalling, regulation of gene expression

## Abstract

Plants align flowering with optimal seasonal conditions to increase reproductive success. This process depends on modulating signalling pathways that respond to diverse environmental and hormonal inputs, thereby regulating the transition to flowering at the shoot apical meristem. In *Arabidopsis*, long-day photoperiods (LDs) stimulate the transcription of *FLOWERING LOCUS T* (*FT*), encoding the main florigenic signal. *FT* activation is mediated by the transcriptional regulator CONSTANS (CO), which binds to the CO responsive elements (COREs) located in the proximal *FT* promoter region. The phytohormone abscisic acid also (ABA) contributes to *FT* activation together with GIGANTEA (GI) to regulate drought escape (DE). Whether CO is a target of ABA and GI actions for the regulation of *FT* is, however, unknown. Here, we report that ABA and its signalling components promote CO recruitment to the CORE1/2, without causing reductions in the diel pattern of CO protein accumulation. ChIPseq analyses show that ABA broadly shapes the CO DNA binding landscape, which is enriched at the promoters of genes involved in the response to abiotic stress. We also found that GI promotes CO recruitment to the CORE1/2 region, and that CO recruitment is required for the accumulation of RNAPol II at the transcription start site of *FT*. Finally, we show that GI and ABA signalling pathways are largely epistatic in the control of flowering time, suggesting their involvement in the same molecular process. Taken together, these observations suggest that varying water deficit conditions modulate CO recruitment and *FT* expression, thus dictating DE strategies in *Arabidopsis*.

**Highlight:** ABA and GIGANTEA signalling promote *FLOWERING LOCUS T* (*FT*) transcriptional activation by regulating the binding of the transcription factor CONSTANS to the proximal *FT* promoter.

## INTRODUCTION

During floral transition plants shift from vegetative to reproductive growth. This complex process is orchestrated by a combination of internal (e.g., age) and external (environmental) signals that ultimately cause transcriptional reprogramming and changes of cell fates in young primordia located at the shoot apical meristem. At the molecular level, these signals are relayed through a multitude of molecular pathways, hormonal signalling cascades and transcriptional events that unfold at different temporal and tissue scales. Variations in daylengths (photoperiod) play a key role in flowering time regulation and its measurement occurs in the vascular tissue of leaves (GARNER And ALLARD, 1920; Wigge et al., 2005; Song et al., 2015; Gendron and Staiger, 2023). Following perception of an inductive photoperiod condition, a specialised protein signal referred to as florigen is produced and exported systemically to activate floral transition at the shoot apical meristem (Kardailsky *et al*., 1999; Kobayashi *et al*., 1999). In *Arabidopsis* the photoperiodic cascade is activated under long day conditions (LDs) and requires the activity of three key genes including *GIGANTEA* (*GI*), *CONSTANS* (*CO*) and *FLOWERING LOCUS T* (*FT*) (Song *et al*., 2015). GI is a key activator of the network and encodes a plant-specific protein with unclear molecular functions. GI forms an oscillatory (diel regulated) complex with blue light receptors to set the stage for the transcriptional activation of *CO* during late afternoon in LDs (Suárez-López *et al*., 2001; Imaizumi *et al*., 2005; Sawa *et al*., 2007; Fornara *et al*., 2009; Ito *et al*., 2012). Light-stabilized CO protein promotes the transcriptional activation of *FT*, encoding the main florigen of *Arabidopsis*, so that peak *FT* transcript accumulation occurs at dusk under standard growth conditions (Kardailsky *et al*., 1999; Kobayashi *et al*., 1999; Valverde *et al*., 2004; Corbesier *et al*., 2007; Song *et al*., 2018).

The molecular mechanisms that lead to the stabilization of CO, its post-translational modifications, binding to specific DNA cis-elements, and interactions with a diverse array of co-regulators are collectively important to finely tune the expression of *FT* (Takagi *et al*., 2023). CO binds to *FT* regulatory regions at the CO Response Elements (*CORE*s) and *P1/2* sites. These are 4-bp TGTG core motifs located in the proximal promoter of *FT* (–285bp from ATG), broadly corresponding to Block A regulatory region (Adrian *et al*., 2010; Tiwari *et al*., 2010; Hayama *et al*., 2017; Lv *et al*., 2021). CO binding to DNA occurs through the positively charged CCT domain, located at the C terminus, which is evolutionarily related to the DNA binding domain observed in NUCLEAR FACTOR-Y subunit A (NF–YA) encoding genes (Wenkel *et al*., 2006). CO forms a trimeric complex with NF–YB and NF–YC subunits to bind to DNA (Ben-Naim *et al*., 2006; Tiwari *et al*., 2010; Gnesutta *et al*., 2017). At the same time, through the N terminal B-BOX domain CO can form multimers, each one being able to form NF–YB/YC complexes via the CTT for high affinity binding to the TGTG core motifs (Lv *et al*., 2021; Zeng *et al*., 2022). Besides NF-YB/C, other mechanisms regulate CO recruitment to chromatin (Bu *et al*., 2014; Wang *et al*., 2016). All in all, these factors together with CO participate in forming local chromatin looping and initiate *FT* transcription (Adrian *et al*., 2010; Tiwari *et al*., 2010; Farrona *et al*., 2011; Cao *et al*., 2014; Tian *et al*., 2021).

Several environmental and hormonal pathways influence CO function and, ultimately *FT* transcriptional activation (Pin and Nilsson, 2012; Conti, 2017). The phytohormone abscisic acid (ABA) is emerging regulator of flowering time, influencing the expression multiple floral targets (Martignago *et al*., 2020). The ABA biosynthesis pathway has been well characterised (Seo and Marion-Poll, 2019) with key steps occurring in the plastids catalysed by the zeaxanthin epoxidase (ABA1) and the nine-cis-epoxycarotenoid dioxygenases (NCEDs) and the cytosol, operated by ABA2 and other enzymes. The ABA signalling cascade comprises a family of soluble receptors known as PYR/PYL/RCARs (Cutler *et al*., 2010) that can bind to ABA. Upon binding the receptor can interact with a clade of protein phosphatases, the PP2C (Fujii *et al*., 2009). This interaction inhibits the phosphatase activity of PP2Cs, allowing their substrate, protein kinases of the SNF1-related protein kinase 2 (SnRK2) group, to auto phosphorylate. Active SnRK2s can in turn phosphorylate downstream components of ABA signalling including the ABA-regulated bZIPs (ABRE binding factors, ABFs) which mediate transcriptional reprogramming (Furihata *et al*., 2006; YOSHIDA *et al*., 2015).

Much of our knowledge about the impact of ABA in flowering time derives from the study of the drought escape response (DE), an acceleration of flowering triggered by water deficit. Under these conditions, water deficit trigger the expression of *FT* and the flowering time regulator *SUPPRESSOR OF OVEREXPRESSION OF CONSTANS 1* (*SOC1*) in leaves, in an ABA-dependent manner and likely through different mechanisms (Riboni *et al*., 2013; Hwang *et al*., 2019). The ABFs in association with NF-YC subunits activate *SOC1* (Hwang *et al*., 2019), whereas little is known about the regulatory role of ABA on CO upstream of *FT*. Recently, the SNF1-related protein kinase 2 (SnRK2) substrate 1 (SNS1) was shown to promote DE by inducing *FT*. SNS1 activity is regulated by phosphorylation and subsequent stabilization by ABA stimulated SnRK2 protein kinases (Katagiri *et al*., 2024). Impairing ABA signalling causes a reduction in *FT* expression with relatively weaker changes in *CO* transcript accumulation suggesting that ABA signalling may act at least in part downstream of *CO* transcriptional activation (Riboni *et al*., 2016). In this process, GI function is key for the ABA – dependent activation of *FT* and DE (Riboni *et al*., 2016). GI might act as a general regulator of ABA signals for its involvement in modulation of physiological traits related to gas exchange (which are also ABA regulated) as well as ABA production (Ando *et al*., 2013; Baek *et al*., 2020; Simon *et al*., 2020). However, *gi* plants also display a general hyper activation of ABA signalling responses, suggesting that the mode of action of GI in DE regulation may be separate (and more general) from its role in regulating ABA signals (Siemiatkowska *et al*., 2022).

In this study tests were made to understand how changes in ABA accumulation could alter *FT* expression through CO. We reveal that CO recruitment to the *FT* promoter is ABA regulated which impacts the recruitment of the RNAPol II. Surprisingly GI is also required for proper CO recruitment to the *FT* promoter, indicating that GI and ABA may operate in the same genetic pathway. Overall, we hypothesise that GI could be an important factor bridging photoperiodic stimuli with ABA signalling to relay water deficit information and regulate flowering time.

## MATERIAL AND METHODS

### Plant material, genotyping, and growing conditions

*Arabidopsis thaliana* plants used in this study are of ecotype Columbia (Col-0) or Landsberg *erecta* (L.*er*). Mutant and transgenic lines of *aba1-6* (Niyogi *et al*., 1998)*, aba2-1* (Leon-Kloosterziel *et al*., 1996)*, nced3/5* (Frey *et al*., 2012)*, abi1-1* (Bertauche *et al*., 1996)*, SUC2::HA:CO* (Jang *et al*., 2009)*, co-10* (Laubinger *et al*., 2006)*, gi-2* (Rédei, 1962)*, gi-100* (Huq *et al*., 2000) were previously described. Seeds were stratified and plants grown under long day conditions (LDs, 16 h light / 8 h dark), under controlled-environment cabinets, as previously described (Riboni *et al*., 2016). Temperature range in the growth chamber fluctuated between 24 °C during the day and 21 °C in the night and air humidity was about 60%. For *in vitro Arabidopsis* growth Murashige and Skoog (MS) medium was prepared by dissolving the MS salt mix (Duchefa) in distilled water. After adjusting the pH solution to 5.8, agar (Duchefa) was added to a final concentration of 0.8% w/v. Sterilized seeds (70% v/v ethanol and 1% w/v Sodium Dodecyl Sulphate, SDS for 10 minutes) were spread onto solidified agar plates, stratified for 2-3 days (4°C and dark) and then moved to the growth chamber. ABA application experiments on soil were performed following the procedure detailed previously (Riboni *et al*., 2016). A mock solution (0.025% v/v ethanol) was used as a mock control. Mutant combinations were selected via PCR of genomic DNA with primers described previously (Kaiserli *et al*., 2015; Riboni *et al*., 2016) or in Supplementary Table 1 and by exploiting markers encoded by the different transgenes; *gi-100, SUC2::CO:CITRINE* (Basta resistance), *SUC2::HA:CO* (GFP expression in seeds).

### Molecular cloning

Cloning was done with Gateway and Multi-Site Gateway (three-fragment vector) cloning technology (Invitrogen). To generate entry clones (pDONR207, Invitrogen), the *AttB1*/*AttB2*-containing sequences were fused to the *CONSTANS* specific primers (Supplementary Table 2). The Phusion High Fidelity DNA polymerase (New England Biolabs) was used for all the PCR reactions. The 5’ and 3’ elements and destination vectors were previously described; *SUC2 promoter / pDONR221 P4-P1r* (Marquès-Bueno *et al*., 2016), *mCITRINE / pDONR221 P2r-P3* (Jaillais *et al*., 2011) and *pB7m34GW* (Karimi *et al*., 2005). All the recombinant destination vectors were transformed into *Agrobacterium* cells, strain GV3101 (Koncz and Schell, 1986), for *Arabidopsis* stable transformation.

The reporter plasmid used for monitoring luciferase (Luc) activity carries a 5’ *FT* regulatory region (1666 bp upstream of ATG). The sequence was custom synthetized (Officinae Bio, Italy) and cloned in the reporter plasmid *pGreenII 0800-LUC* (Hellens *et al*., 2005). This plasmid also contains the *35S:Renilla* (Ren) cassette. The effector construct (*35S:CO*) was built by recombining a CO entry vector (which includes the stop codon) into the Gateway destination vector *p2GW7* (Karimi *et al*., 2002).

### Plants transformation and BASTA selection

Destination vectors *SUC2::CO:CITRINE* and *pSUC2::GW* (here used as a control for Basta resistance) (An *et al*., 2004), were introduced into *Agrobacterium* by electroporation. Transformed *Agrobacterium* were used to generate *Arabidopsis* transgenic plants via the floral dip technique (Clough and Bent, 1998). Transgenic plants were selected on soil after continuous Basta applications (12 mg/l dilution, Agrevo) and single locus insertion events were selected based on a Mendellian 3:1 ratio in T_2_ generation. T_3_ homozygous lines were selected on soil or MS+Basta plates, according to the absence of Basta resistance segregation. Transgenic *SUC2* plants propagated up to the 5^th^ generation by selfing did not show obvious signs of silencing.

### Protoplast Isolation and transactivation Assays

Leaf protoplasts were isolated from 3-week-old Arabidopsis plants as previously described (Iacopino *et al*., 2019). Luc and Ren activities were measured by providing the specific substrates (Dual-Luciferase reporter assay system, Promega) and luminescence was measured with the EnSight plate reader (Perkin Elmer). In each assay, data were expressed as Luc activity relative to Ren.

### Flowering time measurement

Flowering time was measured by scoring the number of rosette leaves, excluding cotyledons. When indicated, flowering was expressed as days to bolting (days between sowing and a visible emergence of the primary inflorescence). Data regarding cauline leaf number (I1 phase) were also recorded for each genotype. To avoid flowering time alterations associated to soil transfer from plates, Basta resistance selection in T1 generation was performed directly on soil.

### RNA extraction and Real-Time qPCR

Total RNA was extracted with QIAzol reagent (Invitrogen) and suspended in RNase-free milliQ dH_2_O. RNA concentration was measured with a UV spectrophotometer and 1000 ng aliquots were used for cDNA synthesis with the Maxima Reverse Transcriptase kit (Thermofisher). Quantitative Real-Time PCR (qPCR) was performed as previously detailed including primers to amplify *CO*, *FT*, *ACT2* and *IPP2* were described (Kaiserli *et al*., 2015; Riboni *et al*., 2016). Primers to amplify *CITRINE* are listed in Supplementary Table 1.

### Nuclei isolation and CO protein detection in *Arabidopsis*

Approximately 100 mg of *Arabidopsis* seedlings were harvested and immediately frozen in liquid nitrogen. Tissue samples were ground by shaking with glass beads in a TissueLyser II (Qiagen, 2 pulses of 30 s each at 28Hz shaking). Leaf powder was suspended in 1.2 ml of cold nuclear isolation buffer (20 mM Tris-HCl, pH 8.8, 25 mM NaCl, 5 mM MgCl_2_, 30% (v/v) glycerol, 5% (w/v) sucrose, 0.5% (v/v) Triton X-100, 0.08% (v/v) β-mercaptoethanol, 0.2% (v/v) SIGMA plant protease inhibitor, 1mM DTT, 1.3 mM PMSF) (Hayama *et al*., 2017). The samples were filtrated trough two layers of Miracloth (Millipore) and centrifuged at *5,000 x g*, at 4 °C, for 10 minutes. The pellet was washed four times with 1 ml of nuclear isolation buffer and, after each wash, pelleted at 4 °C at decreasing speed: *5,000 x g*, *2,700 x g*, *2,200 x g* and *2,200 x g*, 8 minutes each time. Nuclei were suspended in 30 μl of nuclear isolation buffer, mixed with 10 μl of Laemmli Buffer and heated 10 minutes at 95 °C. Samples were centrifuged for 1 minute at *3,000 x g* to pellet all nuclear membranes, and 20 μl of the supernatants (enriched in soluble nuclear proteins) were loaded onto an 10% SDS-PAGE gel. For total protein detection, tissue from 50 mg seedlings was harvested in liquid Nitrogen and lysed in pre-cooled TissueLyser II (Qiagen) for 3 min at the frequency 30/s. Proteins were isolated by homogenisation in 450 µL sample buffer (100 mM Tris-HCl pH 6.8, 20% (v/v) glycerol, 5% (w/v) SDS, 20 mM DTT, 1mM bromophenol blue, 1× proteinase inhibitor cocktail (Sigma) and 80 µM MG132), boiled at 95 °C for 5 min, centrifugated for 1 min at *12000* x g and loaded onto an 4%-12% Bis-Tris Plus SDS Mini Protein Gel run in 1× Bolt MOPS SDS buffer (Thermo Fisher). Proteins were transferred to a nitrocellulose membrane and incubated with anti-HA-Peroxidase, High Affinity antibody (Roche), anti-Histone H3 (anti H3) or anti-Actin (Agrisera), used for loading control. Chemiluminescent signals were detected through a ChemiDoc Touch Imaging System (Biorad) and quantified with the Image Lab software (Biorad).

### ChIP experiments and sequencing

Chromatin immunoprecipitation assays were performed as described previously with minor modification, starting from 1-2 grams (FW) of seedlings (Perrella *et al*., 2018, 2023). A bioruptor sonicator (Diagenode, B01020001) was used to shear the chromatin using the following settings: 40 Cycles, 30 sec ON, 30 sec OFF at high power. With these conditions the size of sheared DNA is below 100 bp. An anti-RNAPol II antibody (AbCam Ab5408) or GFP-Trap magnetic nanobodies (Chromotek) were used. qPCR was performed at the following cycles: 95°C x 3 min; 95°C x 10 sec, 59.5°C x 30 sec (45 cycles); 65°C x 0.05 sec and 95°C x 30 sec (Melting curve). Relative enrichment was calculated as described previously (Kaiserli *et al*., 2015; Perrella *et al*., 2018). Reactions were performed on four technical replicates and two independent biological experiments. Amplicons of the *CBS*, *CORE1/2*, *CORE3*, *CORE4* and *CORE5* regions of *FT* were obtained with primers previously described (Hayama *et al*., 2017), whereas the *TSS* regions of *FT*, *YUCCA8* and *ACTIN2* were amplified with primers lcm217/lcm218, *YUC8 FW*/*RV* and *ACT2 FW*/*RV*, respectively (Supplementary Table 1).

For ChIP-sequencing 5 grams (FW) of seedlings were used for each genotype/replicate and processed as described above. Inputs and ChIP samples (two biological replicates) were quantified with Qubit dsDNA Quantification High sensitivity Assay Kit (Invitrogen). For library preparation, the VAHTS Universal Plus DNA Library Prep Kit for Illumina (ND617-02, Vazyme, Nanjing, China) was used, following the manufacturer’s recommendations without fragmentation. Library quality control was performed using the QSep-400 (BiOptic, Taipei, China) and Qubit 3.0 (Thermo Fisher Scientific, Wilmington, USA). The qualified library was sequenced on the Illumina Novaseq X platform (Illumina, San Diego, USA) with paired-end 150 bp (PE150) mode. Library construction and sequencing were performed at Biomarker Technologies (BMKGENE) GmbH. Mapping of raw reads and peak calling followed the greenscreen pipeline (Klasfeld *et al*., 2022) with slight modifications. The adapted code is available at https://github.com/beaLando/chipseq_aba.git. In brief, fastp (Chen *et al*., 2018) was used to trim and filter reads after quality checks in fastqc (Andrews S., 2010). Mapping of filtered reads was then executed with Bowtie2 with TAIR10.1 as reference genome. Peak calling was done in macs2 (Gaspar, 2018) for each replicate separately using input samples as background. Peaks found for each replicate were then processed with irreproducible discovery rate (IDR) to retain reproducible peaks and with a global IDR (FDR corrected) greater or equal to –log10(0.1). This list was then further filtered to retain only peaks falling outside greenscreen’s blacklisted regions. Further quality checks were executed with qualimap (García-Alcalde *et al*., 2012) and deeptools (Ramírez *et al*., 2014). Finally, ChIPseeker (Yu *et al*., 2015) was used to estimate how frequently peaks would fall within gene promoters or other genomic regions.

Because the two replicates of *SUC2::CO:CIT aba1-6* resulted in very low mapping rates (between 3-15%; Table S1), we simulated a peak calling analysis on *SUC2::CO:CIT*, by subsampling mapped reads until 3-15% mapping rate was achieved. The logic for subsampling *SUC2::CO:CIT* was the following: if a sizeable number of peaks could still be called for *SUC2::CO:CIT* at low mapping rates, the low number of called peaks in the *SUC2::CO:CIT aba1-6* mutant could be due to a real biological effects leading to low binding. The ChIPseq data from this study is available under the GEO accession number GSE291256.

### Metanalysis studies of RNA-seq and ChIP-seq datasets

We retrieved CO ChIPseq peaks data (de los Reyes *et al*., 2024) – GEO accession number GSE222657 – and gene expression data of upregulated DEGs in *SUC2::CO* and *35S::CO* genotypes relative to the WT (same study). Using ChIPseeker, CO peaks were then annotated with gene IDs based on the genes’ distance from the peak. Using GeneOverlap (Shen, 2024) we assessed the overlap in Click or tap here to enter text.peaks, or genes in proximity of peaks, and their significance, for the two datasets – our study and (de los Reyes *et al*., 2024) using the number of protein-coding genes in *A. thaliana* as background. We then obtained the intersection between genes falling next to *SUC2::CO:CIT* peaks and upregulated *SUC2::CO* DEGs compared to WT. Based on this intersection, we ran an enrichment analysis for biological process terms using the union of all *SUC2::CO* upregulated DEGs and genes falling next to *SUC2::CO:CIT* peaks as background. The same procedure was followed for *35S::CO* peaks and DEGs. All plots were produced with ggplot2 (Wickham, 2011) and Inkscape.

### Statistical analysis

Analysis of variance (ANOVA) was conducted with linear models where flowering time/gene expression was predicted by genotype according to the formula: flowering time (gene expression) ∼ genotype. Here, “flowering time” was expressed as either one of: leaf number at bolting, days to bolting, or number of l1 nodes. “genotype” refers to the allelic status at *ABA1*/*2* (WT, *aba1-6*, *aba2-1*) or *GI* (WT, *gi-2*, *gi-100*) loci, and different isogenic lines for the various mutants. When *aba1* mutants were in different backgrounds (WT, *SUC2::CO:CIT*, *SUC2::HA:CO*), the same model was run separately for the different backgrounds and the models were qualitatively compared. In Fig. 2A, the model used was gene expression ∼ genotype*timepoint. If the interaction was not statistically significant, only the additive model (genotype + timepoint) was considered for post-hoc tests.

**Figure 1.**
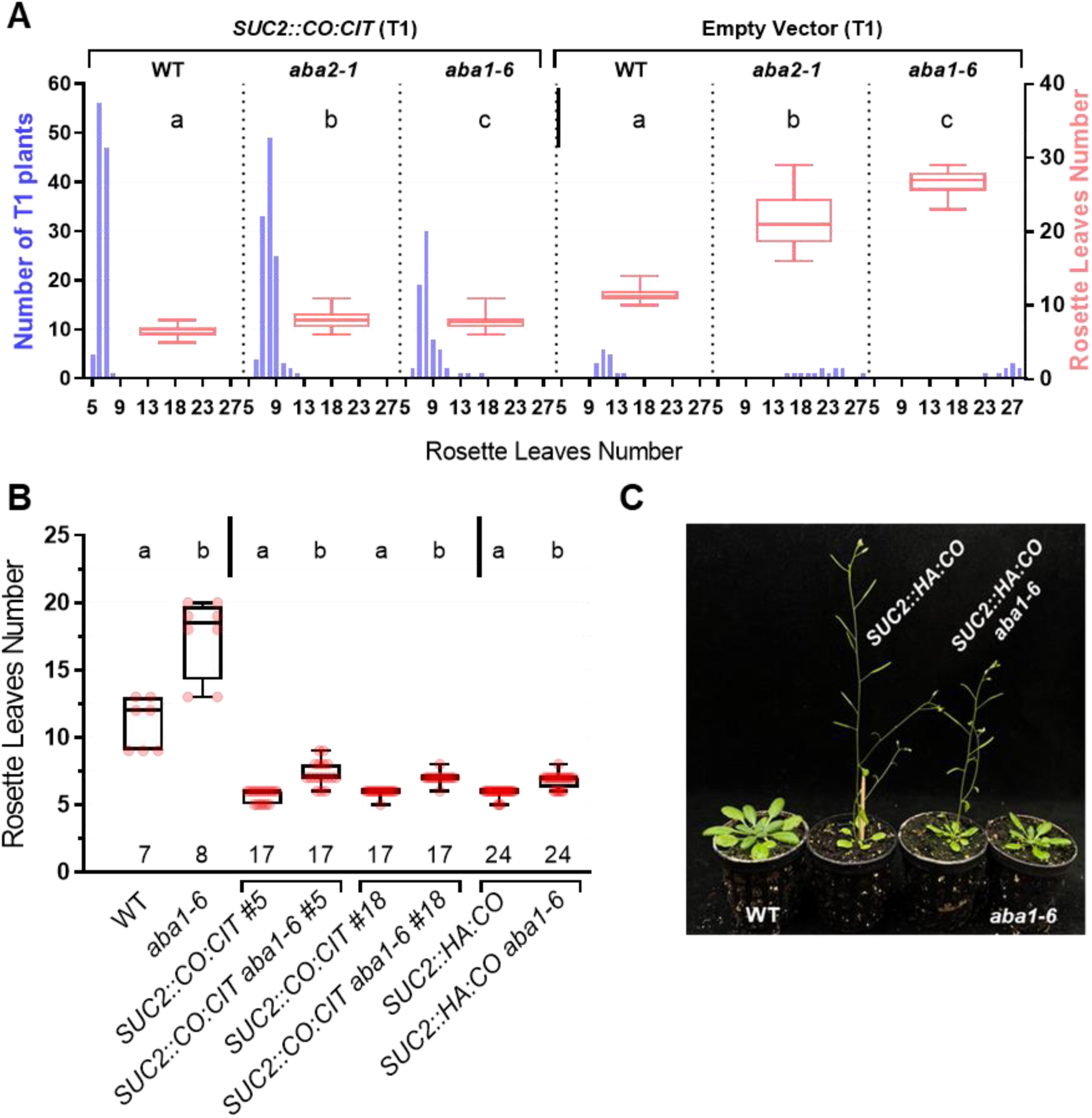
ABA promotes CO function in the phloem companion cells. (A) Boxplot of flowering time (right Y axis, rosette leaves number) of Basta resistant transgenic T1 plants of the indicated genotypes. Whiskers represent the range, min to max values. The left Y axis shows the number of plants scored for each genotype, and their phenotype distribution according to rosette leaves number (X axis). An ANOVA test to assess the impact of *aba1* and *2* mutations was run separately for *SUC2::CO:CIT* and empty vector – transformed plants (vertical bar). For both groups (empty vector or *SUC2::HA:CO* transformed), the genotype was a statistically significant predictor of leaf number at bolting (*p* < 0.001). Letters at the top of boxplots indicate statistically significant differences (*p* < 0.05) according to a Tukey post-hoc test. (B) Boxplot of flowering time of the indicated genotypes and isogenic lines derived from the introgression of different *SUC2::CO* transgenes (wild type Col-0) into *aba1-6*. An ANOVA test to assess the impact of *aba1-6* mutation was run separately for transgenic (*SUC2::CO:CIT/SUC2::HA:CO*) and non-transgenic controls (vertical bar). For both groups, the *aba1-6* genotype was a statistically significant predictor of leaf number at bolting (*p* < 0.001). Letters at the top of boxplots indicate if genotypes showed statistically significant differences (*p* < 0.05) according to a Tukey post-hoc test. The number of samples analysed for each genotype is shown at the bottom of the graph (C) Representative plants of the indicated genotypes pictured 27 days after sowing.

**Figure 2.**
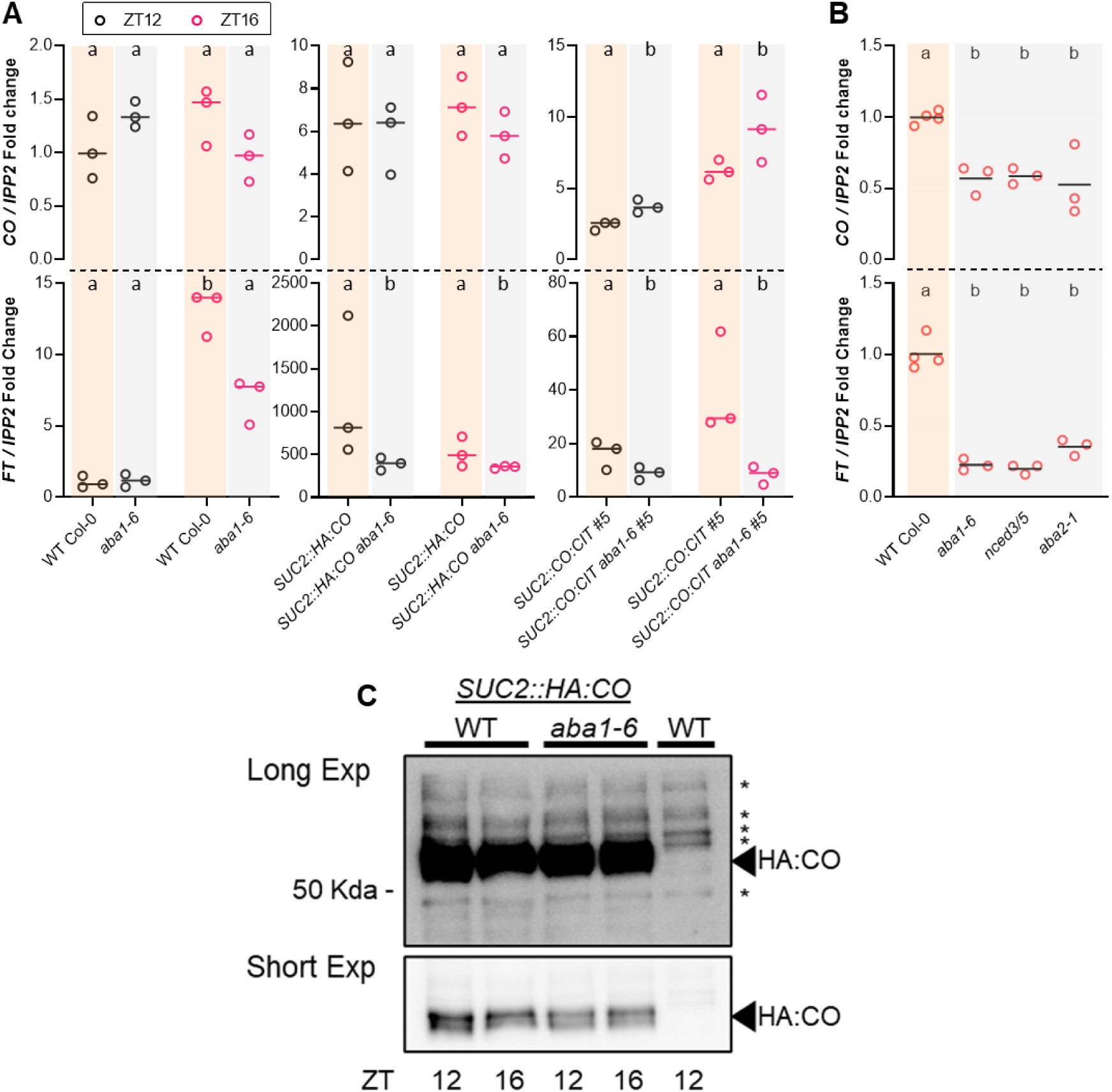
ABA regulates CO function without altering its accumulation in *Arabidopsis* (A) Real-Time qPCR of *CO* (upper panels) and *FT* (lower panels) transcripts accumulation in seedlings of the indicated genotypes grown on plates for 13 days and harvested at the indicated Zeitgeber times (ZT, h) in the diurnal phase. Horizontal Lines represent the mean fold change of *IPP2*-normalised *CO*/*FT* accumulation relative to the wild type (the three biological replicates are shown as dots). Each replicate comprises approximately 50 mg of seedlings. Different letters denote differences (*p* < 0.05) between genotypes within each ZT according to post-hoc tests. (B) Same as A, from seedlings grown for 9 days under LDs and harvested at ZT16. Different letters denote different (*p* < 0.05) groups according to post-hoc tests. In (A) and (B), the plot background colours denote the ABA producing vs the ABA deficient genotype (C) Immunoblot detection of HA:CO protein from nuclear fractions of seedlings grown on plates for 12 days under LDs and harvested at the indicated ZT points. WT, non-transgenic Col-0 plants were used as negative controls whereas asterisks denote non-specific bands detected by the antibody and used as loading controls. Two exposures of the same blots are shown. Numbers refer to molecular mass according to the migration of a protein ladder.

Before running the ANOVAs, the condition of homogeneity of variances between groups was checked with a Bartlett’s test in base R (R Core Team 2023). If group variances were not homogeneous, a log transformation was applied to the data. All ANOVA models were conducted with the R function lm (R Core Team 2023) using sum contrasts and significance of predictors was assessed with the ANOVA function from the car package (Fox and Weisberg, 2018). Post-hoc Tukey tests to assess differences between each pair of genotypes were conducted with the R packages multcomp (Hothorn *et al*., 2008) and emmeans (Lenth, 2023). Even for models including timepoint as predictor, post-hoc tests were conducted exclusively to assess differences between genotypes within timepoints.

## RESULTS

### CO function is sensitive to ABA signalling status in the phloem companion cells

To evaluate the contribution of ABA accumulation in regulating CO function and *FT* expression we generated transgenic plants constitutively expressing *CO* in different ABA-impaired/defective backgrounds. Since CO regulates *FT* in the phloem companion cells (An *et al*., 2004) we engineered a *CO:CITRINE* (*CO:CIT*) translational fusion gene under the control of the phloem companion cell-specific *SUCROSE-PROTON SYMPORTER 2* (*SUC2*) promoter (Truernit and Sauer, 1995). Basta resistant T1 plants of wild-type (Col-0) and two different ABA-deficient backgrounds (*aba1-6* and *aba2-1*, impaired at different steps of the ABA biosynthetic pathway) (Nambara and Marion-Poll, 2005) displayed a clear early flowering phenotype compared with the corresponding genotypes transformed with an empty vector (Fig. 1A). *SUC2::CO:CIT aba1-6* and *aba2-1* plants showed a mild but significant delay in flowering (*p* = 6.1E-07 and 1.2E-14, respectively) compared with the wild type transformed with *SUC2::CO:CIT*. Ruling out an indirect effect of reduced ABA production on *SUC2* promoter activity, we detected similar levels of *CIT* transcript accumulation across randomly chosen T1 plants (Supplementary Figure S1). This suggests a role for ABA production downstream of *CO* transcriptional activation. In support of this view, we observed similarly reduced level of *FT* accumulation in *SUC2::CO:CIT* in either *aba1-6 or aba2-1* independent T1 transgenic events compared with *SUC2::CO:CIT* transgenics events in the wild-type background (*p* = 0.047 and 0.026, respectively, Supplementary Figure S1). *FT* transcript levels detected in *SUC2::CO:CIT aba2-1/aba1-6* plants were, however, slightly increased compared with the wild type transformed with an empty vector, consistent with their earlier flowering.

To confirm the observed flowering time phenotypes we crossed two homozygous T3 *SUC2::CO:CIT* plants with *aba1-6* to generate isogenic lines (#5 and #18). We also performed genetic crosses between *SUC2::HA:CO* plants (Jang *et al*., 2009) and *aba1-6* to obtain *SUC2::HA:CO aba1-6*. *SUC2::HA:CO* plants were extremely early flowering, similar to *SUC2::CO:CIT* plants. However, in the *aba1-6* background, the early flowering phenotype conferred by *SUC2::HA:CO* or *SUC2::CO:CIT* was significantly alleviated (*p* < 4.6E-06, Fig. 1B and 1C). Collectively, our findings suggest that ABA may promote CO function and *FT* transcript accumulation to activate flowering.

### Reduced ABA levels impair *FT* transcript accumulation

We next aimed to evaluate potential modes of regulations of CO function controlled by ABA. Using stable transgenic isogenic lines expressing *SUC2::HA:CO* or *SUC2::CO:CIT #5* in the wild-type or *aba1-6* backgrounds, we confirmed that *FT* transcript accumulation was reduced in *aba1-6* plants compared with the corresponding ABA producing WT background (*p* < 3.33E-03, Fig. 2A) irrespective of the time point analysed (ZT12 and 16 under LDs). In contrast, *CO* levels remained not affected in *SUC2::HA:CO aba1-6* plants or even increased (*p* = 1.82E-03) in *SUC2::CO:CIT#5 aba1-6* compared with the corresponding *SUC2::CO:CIT #5*. Interestingly, we also detected reduced levels of *FT* transcript accumulation in *aba1-6* plants compared with the wild type (*p* = 1.92E-04) at ZT16 when we also observed a marginal reduction (*p* = 0.06) in *CO* transcript levels. To investigate further the role of ABA production in *FT* transcript accumulation we analysed 9-day-old seedlings of wild type and strong ABA deficient mutant backgrounds, including *aba1-6*, *aba2-1* and double mutants of *9-cis-epoxycarotenoid dioxygenase* (*nced*) *3* and *5* (Frey *et al*., 2012). Compared with the wild type, at ZT16 we observed reduced *FT* accumulation in all ABA deficient mutants (*p* < 6.09E-06) and correspondingly reduced levels of *CO* transcript (*p* < 0.01) (Fig. 2B). Collectively, our data indicate a role for ABA in promoting *FT* accumulation with ABA affecting both *CO* expression and its function.

To gain further insights into the potential mode of ABA-dependent stimulation of CO we analysed HA:CO protein levels derived from *SUC2::HA:CO* and *SUC2::HA:CO aba1-6* plants. HA:CO protein accumulation was broadly similar between the two genotypes at ZT12 and ZT16, suggesting that reduced ABA accumulation impairs HA:CO protein function but not HA:CO accumulation (Fig. 2C). We next extended this analysis to morning timepoints including ZT 0.5 (i.e., within 30 min after the switch on of the lights), ZT4 till midday-dusk (ZT8 and ZT12). At ZT0.5 we observed a much stronger increase in HA:CO accumulation in *SUC2::HA:CO aba1-6* plants compared to *SUC2::HA:CO* (Supplementary Figure S2). This peak of accumulation was, however, transient and might suggest a role of ABA in repressing this phase of CO stabilization in this early morning timepoint (Valverde *et al*., 2004; Song *et al*., 2012; Hayama *et al*., 2017). HA:CO protein was still clearly detectable in all the other time points in *SUC2::HA:CO aba1-6* plants further pointing to defective CO function with respect to *FT* and consequent flowering activation.

To independently test for potential effects of ABA on HA:CO protein abundance we exogenously applied three different concentrations of ABA (0.25 –2.5 –25 µM) on a daily basis for 12 days (Riboni *et al*., 2016). Tissue from ABA-treated plants was collected at ZT4 and ZT12 and HA:CO protein was detected from nuclear extracts. We did not observe obvious changes in HA:CO protein levels at any of the concentrations and time points assayed (Supplementary Figure S3). In addition, we did not observe obvious changes in the pattern of HA:CO protein gel mobility in the ABA deficient background *aba1-6* (Supplementary Figure S3), which are determined by its phosphorylation status (Sarid-Krebs *et al*., 2015; Chen *et al*., 2020). In summary, our data indicate that CO function might be stimulated by ABA in the cells where *FT* is transcriptionally activated without clear changes at the post translational level.

### ABA stimulates the recruiting of CO to the *FT* promoter region

We next investigated what other aspects of CO function could be influenced by ABA status. We first set up a transient reporter expression assay in *Arabidopsis* mesophyll protoplasts, by fusing a 1666 bp promoter region of *FT* upstream of the *LUCIFERASE*-encoding sequence (hereafter *FT_pro_::LUC*). This promoter region encompasses several cis elements that are critical for the *FT* transcriptional activation, including the well characterised CO responsive element (*CORE*)*1/2* (Tiwari *et al*., 2010). Co-transformation of *FT_pro_::LUC* with *35S::CO* vectors resulted in a strong activation of LUC activity, indicating that the fused promoter can respond to CO binding (Supplementary Figure S4). Co-incubation of *FT_pro_::LUC/35S::CO* transformed protoplasts in presence of ABA (10 and 100 nM), however, did not produce significant changes in LUC activity suggesting that exogenous ABA does not directly influence CO-mediated stimulation of the *FT* promoter, at least in the absence of a native chromatin context.

We therefore evaluated CO recruitment to the *FT* promoter through a Chromatin Immunoprecipitation (ChIP) assay by comparing CO binding in isogenic transgenic seedlings of *SUC2::CO:CIT* #5 (wild type and *aba1-6* backgrounds) under LDs. Consistent with previous reports (Cao *et al*., 2014; Song *et al*., 2016; Hayama *et al*., 2017; de los Reyes *et al*., 2024), at ZT12 we observed a strong enrichment of the CO:CIT protein at the *CORE1/2 FT* promoter regions (Fig. 3 and Supplementary Figure S5). In contrast, such CO:CIT enrichment was much decreased (approximately 4X) in the *aba1-6* background, suggesting that ABA promotes CO recruitment to the *CORE1/2* region. A much weaker CO:CIT binding was also observed at the *CORE3* position (Hayama *et al*., 2017), located in the second intron of *FT* and the *CBS* (*CAAT Binding Site*) region, located at 1.8kb upstream of the ATG start codon – Block B enhancer – (Adrian *et al*., 2010). These enrichment signals were also reduced in in the *aba1-6* background.

**Figure 3.**
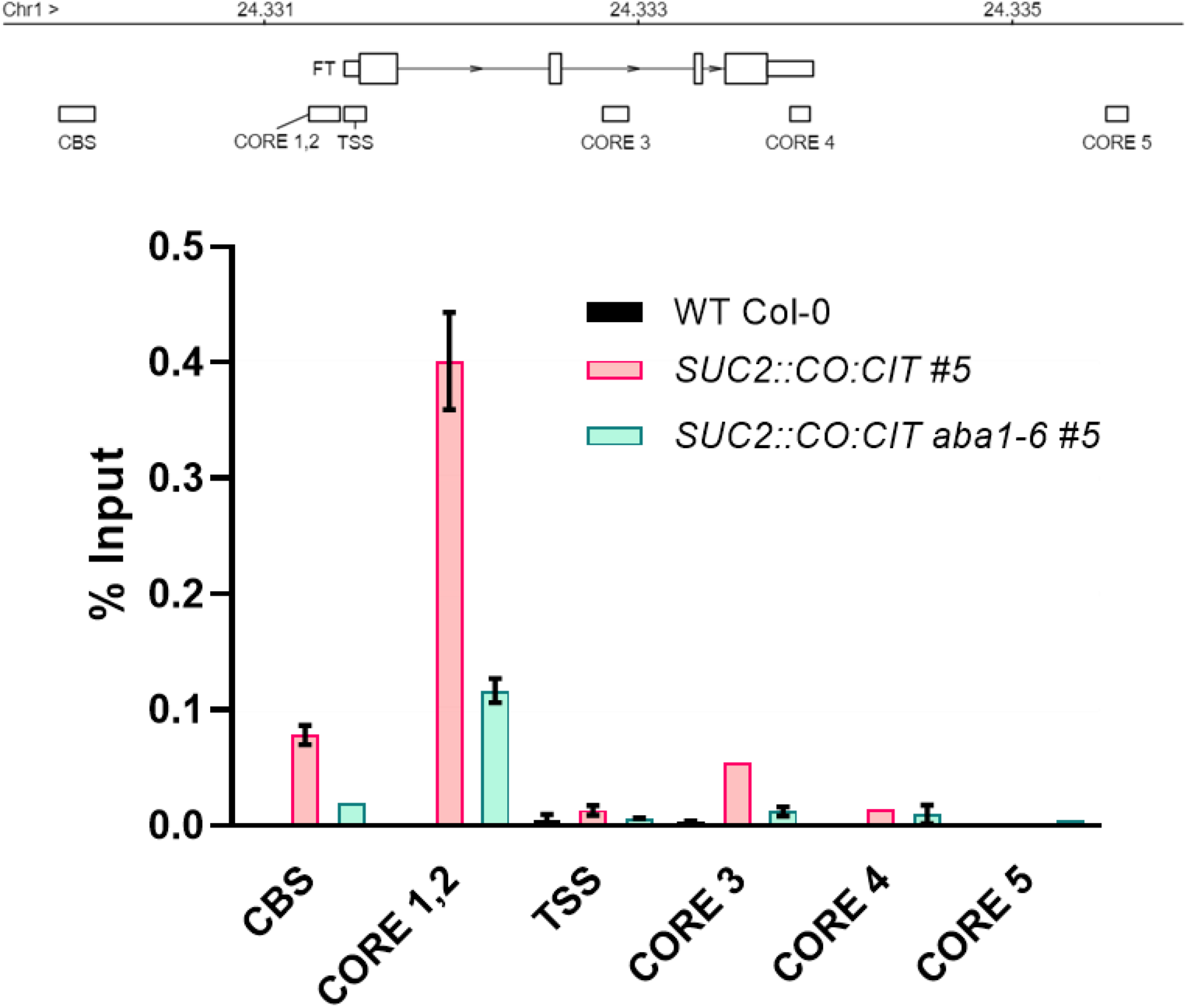
ABA promotes CO recruitment to the *FT* promoter Schematic representation of the *FT* locus (upper panel). Numbers indicate the genomic coordinates (Mbp) of the *FT* region. The diagram illustrates the *FT* transcript, including the 5’ and 3’ untranslated regions (UTRs) (small rectangles), exons (large rectangles), and introns (lines). The location of the sequences amplified by Real-Time qPCR – the CO responsive elements (CORE 1,2,3,4,5), the transcription start site (TSS) and the CAAT binding site (CBS) – are shown below as rectangles. The bottom panel shows the relative enrichment of CO:CIT on the *FT* promoter in isogenic lines *SUC2::CO:CIT #5* (WT and *aba1-6* mutant backgrounds) grown on plates and harvested at ZT12. Non transgenic, WT Col-0 was used as negative control. Bars are the mean ± SE (n = 4 technical replicates). An independent experiment is shown as supplementary material.

To gain a better resolution and coverage of the CO:CIT DNA binding landscape we carried out a ChIP experiment followed by sequencing (ChIPseq). We sequenced the immunoprecipitated DNA derived from *SUC2::CO:CIT*#5 (wild type and *aba1-6* backgrounds) at ZT12 in two biological replicates. We initially retrieved around 20000 peaks per replicate in the *SUC2::CO:CIT* genotype, but only 7312 were retained after irreproducible discovery rate (IDR) filtering (Figure 4A, Table S2 and S3). These peaks were mostly located in proximity of promoter (ca. 60%) and distal-intergenic (ca 32%) regions (Figure 4B) which resulted in 5755 unique genes being recovered in proximity of peaks after annotation (Table S3). The same procedure resulted in around 150-250 peaks called and 1 peak retained in the *SUC2::CO:CIT aba1-6* line, although very low mapping rates (3-15%) were achieved in this background (Figure 4A, Table S2 and S3). We thus performed subsampling of mapped reads in the *SUC2::CO:CIT* genotype and called peaks again at lower mapping rates. Even at mapping rates close to 3-15% (corresponding to 949907 mapped reads per replicate), it was possible to retrieve around 1300 peaks per replicate for *SUC2::CO:CIT*, that is over 1000 more peaks than in the mutant *SUC2::CO:CIT aba1-6* (Table S2). From a qualitative perspective, the mapping density around transcription start sites for all immunoprecipitated replicates showed no enrichment of reads in *SUC2::CO:CIT aba1-6*, while enrichment was visible for *SUC2::CO:CIT* (Figure 4A). This suggests that, despite low DNA yields likely affect our results for the *SUC2::CO:CIT aba1-6* mutant, the almost total absence of reproducible peaks could be partly attributed to low binding by CO:CIT in *aba1-6*.

**Figure 4.**
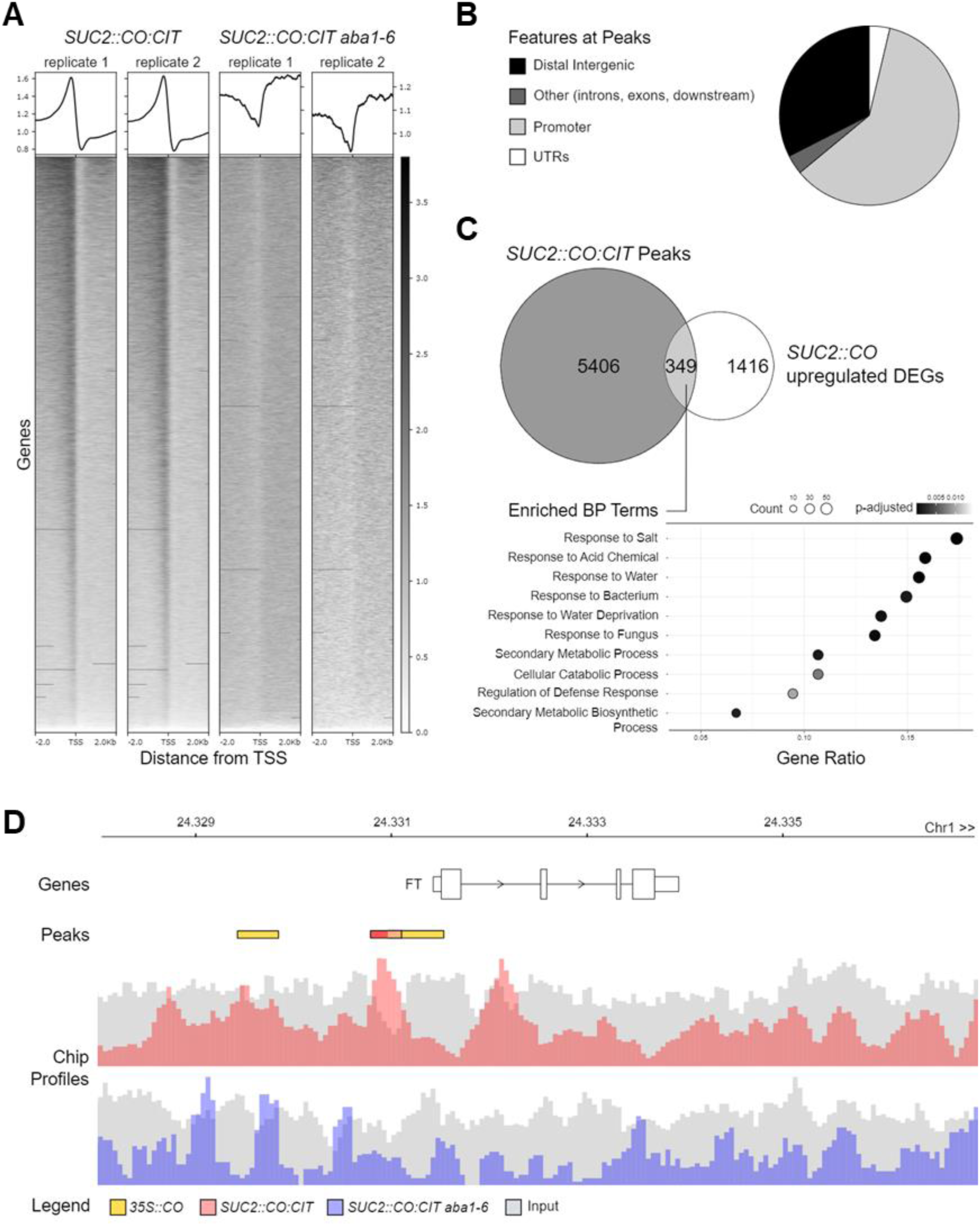
The genome wide CO binding is promoted by ABA. (A) Density of mapped reads around the TSS of known genes following ChIPseq of *SUC2::CO:CIT#5* (WT and *aba1-6* mutant backgrounds) plants (two replicates each) grown on plates and harvested at ZT12. (B) proportion of ChIPseq peaks detected in *SUC2::CO:CIT #5* plants (after filtering by IDR and greenscreen blacklisted regions) falling in proximity of various genomic features. (C) intersection between genes falling next to *SUC2::CO:CIT #5* ChIPseq peaks and *SUC2::CO* upregulated genes, and enrichment of biological process terms for the intersection, where also *FT* is found. (D) Distribution of ChIPseq peaks detected in this study across the *FT* genomic region. The bar charts represent the ChIPseq averaged profiles for *SUC2::CO:CIT #5* (upper panel) and *SUC2::CO:CIT #5 aba1-6* (lower panel) superimposed to their respective inputs. Significant peaks, including those found in the ChIPseq of *35S::CO* plants (de los Reyes *et al*., 2024), are highlighted as colour-coded rectangles.

The overlap of peaks detected in *SUC2::CO:CIT* and those found in *35S::CO* (de los Reyes *et al*., 2024) was not statistically significant (Fisher’s exact test *p* > 0.05, overlap = 252, total = 10272), but the overlap in genes closest to peaks was (*p* < 0.05, overlap = 648, total = 7823). This might be due to differences in how peaks were called and filtered, as well as differences in the promoters used in the two different studies. The comparison of *SUC2::CO:CIT* peaks associated with corresponding upregulated DEGs found in *SUC2::CO* (de los Reyes *et al*., 2024) further disclosed an enrichment for biological process terms related environmental stimuli or stress (abiotic and biotic) responses, including response to salt and water deficit (Fig 4C). A similar over representation of these terms was independently confirmed when comparing CO peaks derived from *35S::CO* and DEGs of *35S::CO* (de los Reyes *et al*., 2024) (Figure S6).

As expected, *FT* was found within the *SUC2::CO:CIT* ChIP peaks – *SUC2::CO* DEGs and *35S::CO* ChIP peaks-DEGs intersections. We found overlapping CO ChIP peaks in the *FT* CORE1/2 region in the two ChIP datasets indicating that the CORE1/2 region represents the main CO binding site *in vivo* (Figure 4D). The ChIPseq analyses also identified potential peaks in regions corresponding to the CBS but not the CORE3. The CBS peak was not significant in our study, but present in *35S::CO* ChIP dataset, indicating weaker binding whereas no binding could be confirmed around the CORE3. Notably, the CORE1/2 peak was absent in *SUC2::CO:CIT aba1-6*, further supporting a role for ABA in promoting CO chromatin binding.

### ABA promotes RNAPol II recruitment to the *FT* promoter

To define the role of ABA in regulating the early transcriptional events at the *FT* promoter we monitored the RNA Polymerase II (RNAPol II) recruitment. We assayed the Ser5 phosphorylated pool of RNAPol II, which in Arabidopsis is distributed over the transcribed region of regulated genes (Obermeyer *et al*., 2023). In the wild type at ZT12 we could detect a peak of RNAPol II enrichment primarily around the *FT* transcription start site (TSS) (Fig. 5A and Supplementary Figure S7). In the *co* and the *gi* mutants, RNAPol II occupancy was much reduced compared with the wild type. This data indicates that the assay is a sensitive readout for CO-stimulated recruitment of RNAPol II to the *FT* promoter (although it does not imply a physical contact between CO and the RNAPol II) (Fig. 5A). Remarkably, we observed a similar strong reduction in RNAPol II occupancy at the TSS in ABA deficient mutants *aba1-6* and *aba2-1* (Fig. 5A). Ruling out a general defect in RNAPol II recruitment at actively transcribed genes in ABA deficient or other mutants utilised in these assays, we observed no clear changes in RNAPol II occupancy at the TSS of *ACTIN2* and *YUCCA8* genes in the same panel of genotypes (Supplementary Figure S8). Therefore, the reduced occupancy of RNAPol II could depend on the lower levels of CO recruitment to the *CORE1/2* region observed in ABA deficient mutants. ABA signals promote the transcriptional activation of *CO* in *Arabidopsis* and other *Brassicaceae* (Yoshida *et al*., 2010; Koops *et al*., 2011; Zhang *et al*., 2022) and we observed reduced *CO* transcript levels in ABA deficient backgrounds, albeit only at the peak of *FT* accumulation (ZT16) (Fig. 2B). To verify further the post-transcriptional control of CO mediated by ABA and its impact on RNAPol II recruitment, we examined the same *FT* regions in *abscisic acid insensitive 1-1* (*abi1-1*) mutants. *abi1-1* encodes a dominant allele of the ABI1 (encoding a PP2C) which severely impairs ABA signalling (Bertauche *et al*., 1996). *abi1-1* plants show similar levels of *CO* transcript accumulation during the day, but diminished levels of *FT* accumulation (Riboni *et al*., 2016). Consistent with this observation, RNAPol II occupancy was reduced in *abi1-1* mutants compared with the wild type (Fig. 5B and Supplementary Figure S9). Interestingly, RNAPol II recruitment in L.*er* (unlike Col-0) was reproducibly enriched primarily towards the *CORE1/2* regions. These results suggest that impairing ABA signalling may also cause similar reductions in CO occupancy at the *FT* promoter which impacts correct RNAPol II recruitment, despite RNAPol II positioning is subject to unknown ecotype-specific controls. Finally, we assessed the role of ABA on RNAPol II recruitment to the *FT* promoter in the presence of constitutive CO accumulation, thus independently of ABA-mediated transcriptional effects on *CO*. We observed a clear decrease in RNAPol II enrichment at the TSS of *FT* in *SUC2::HA:CO aba1-6* compared to *SUC2::HA:CO*, which correlates with the reduced levels of CO occupancy at the *FT* promoter detected in previous ChIP assays (Fig. 5C and Supplementary Figure S7). Notably, RNAPol II was more strongly enriched in *SUC2::HA:CO* plants compared with the wild type, further supporting the limiting role of CO accumulation in RNAPol II recruitment.

**Figure 5.**
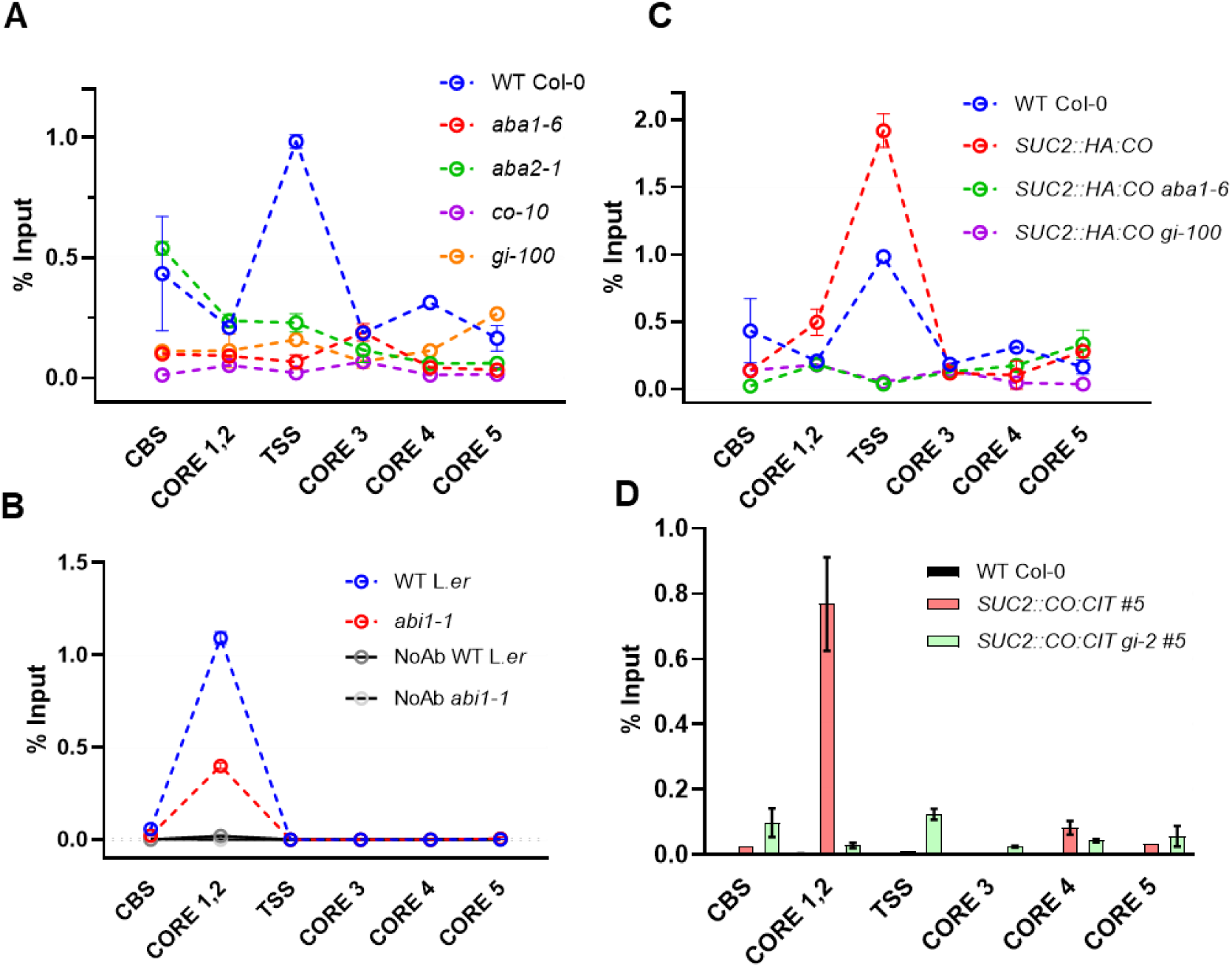
ABA production and signalling are required for proper RNA Pol II recruitment at the TSS of *FT* (A to C) RNAPol II deposition on the *FT* locus in the indicated genotypes expressed as % of input. Each data point represents the mean ± SE (n = 4 technical replicates). Seedlings were grown for 12 days under LDs and harvested at ZT12. In (B), no antibody denotes a negative control for the ChIP assay. In (A) and (C) the WT Col-0 data are the same as samples derived from the same experiment. Iindependent experiments are shown as supplementary material. (D) Relative enrichment of CO:CIT on the *FT* promoter in isogenic lines *SUC2::CO:CIT #5* (WT and *gi-2* mutant backgrounds) at ZT12. Non transgenic, WT Col-0 was used as negative control. Bars are the mean ± SE (n = 4 technical replicates). An independent experiment is shown as supplementary material.

### ABA and GI largely act in the same pathway to control CO recruitment to the *FT* promoter and promote floral transition

Previously we reported the key role for GI in mediating flowering under water deficit conditions (Riboni *et al*., 2016). *gi* mutants are defective in *FT* upregulation under control (Song *et al*., 2014) or water deficit conditions (Riboni *et al*., 2016) even in genetic backgrounds overexpressing *CO*. Consistent with these findings, we observed a strong reduction in RNAPol II recruitment to the TSS of *FT* in *SUC2::HA:CO gi-100* lines, suggesting that GI might also be important for CO recruitment (Fig. 5C and Supplementary Figure S7). A ChIP assay using *SUC2::CO:CIT gi-2* #5 further revealed reduced levels of CO:CIT recruitment to the *CORE1/2* region compared to *SUC2::CO:CIT* #5 (Fig 5D). However, flowering time of *SUC2::CO:CIT gi-2* #5 plants was similar to *SUC2::CO:CIT* #5 indicating that an excess of transgene-derived CO:CIT expression could ultimately compensate for its reduced recruitment observed at the *FT* promoter in *gi* mutants (Supplementary Figure S10).

Because both ABA and GI mediate CO recruitment to the *FT* promoter, we tested whether they acted in the same genetic pathway. We thus compared flowering time (bolting time and leaves counts) of *SUC2::HA:CO aba1-6* and *SUC2::HA:CO gi-100* with triple mutants of *SUC2::HA:CO aba1-6 gi-100* under LDs. As expected, flowering time of *SUC2::HA:CO aba1-6* was significantly delayed compared to *SUC2::HA:CO* plants both in terms of number of rosette leaves (*p* = .01) or bolting time (*p* = 1.78E-11) (Fig. 6A and Supplementary Figure S11). We did not detect a delay of flowering in any *SUC2::HA:CO gi* mutants combination (i.e., *gi-2, gi-100*) compared to *SUC2::HA:CO* in terms of leaf numbers, but we observed a significant delay in days to flowering (*p* = 1.95E-05 for *SUC2::HA:CO gi-2*). Finally, we compared the impact of combined *aba1-6* and *gi-100* mutant alleles in the *SUC2::HA:CO* background. This showed a minor but significant (*p* < 0.01) aggravation of flowering time of *SUC2::HA:CO aba1-6 gi-100* compared with *SUC2::HA:CO aba1-6* in terms of rosette leaves but no consistent effects in days to flowering (Fig. 6A and Supplementary Figure S11), suggesting a partially epistatic relationship between GI and ABA signalling in this process. We also scored the number of I1 coflorescences (i.e., nodes presenting side shoots derived from the main inflorescence, Supplementary Figure S11). I1 nodes were similarly reduced in *SUC2::HA:CO* plants and *SUC2::HA:CO gi* backgrounds. We observed a significant increase in I1 nodes in *SUC2::HA:CO aba1-6* (*p* = 0.002) but no further aggravation of this phenotype in *SUC2::HA:CO aba1-6 gi-100* mutants lines. Therefore, GI and ABA signalling action is largely non additive to regulate flowering time and during inflorescence development in genetic background that overexpress *CO*.

**Figure 6.**
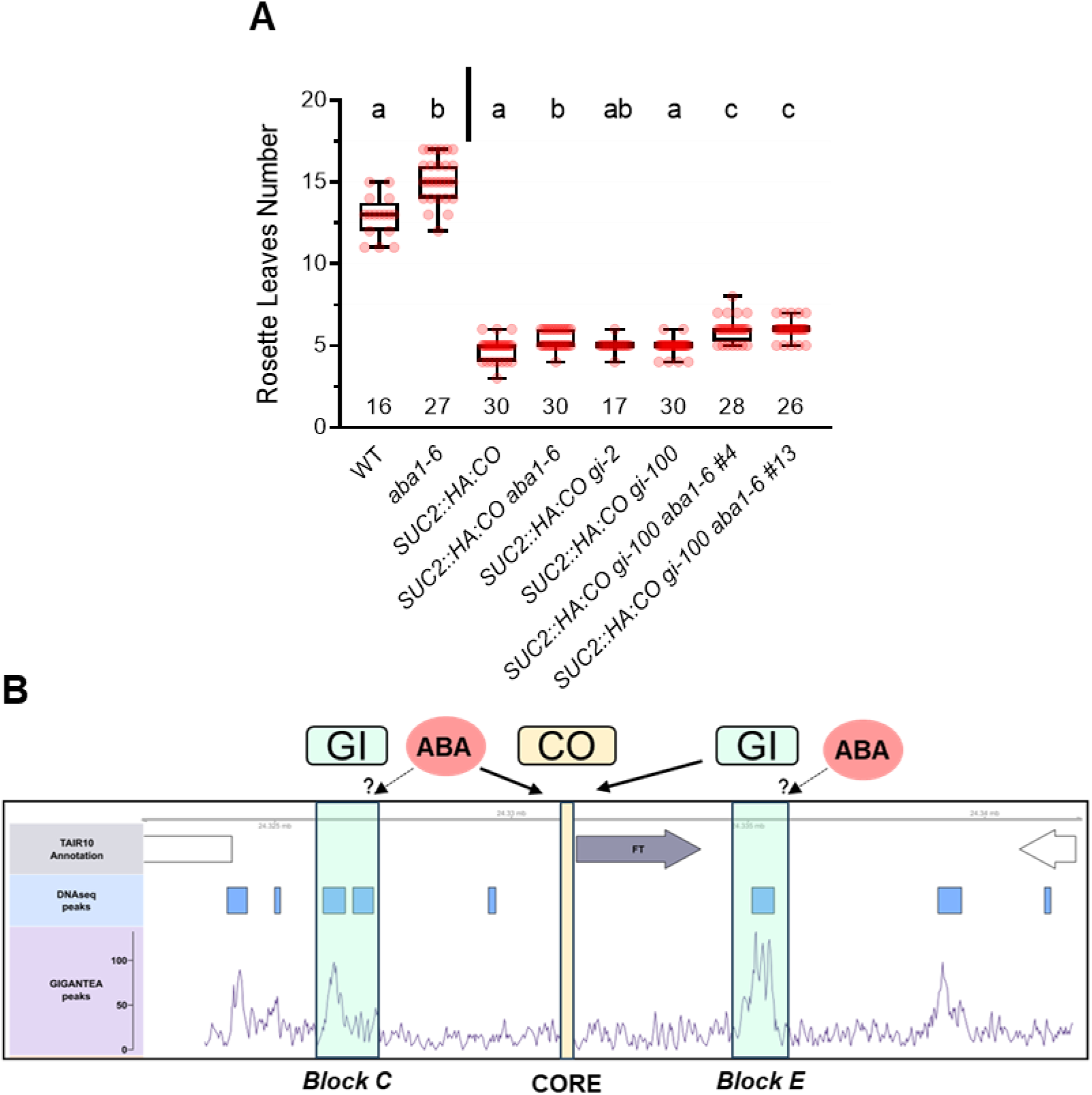
ABA signalling and *GI* act together to promoter CO recruitment to the *FT* promoter and regulate flowering time. (A) Boxplot of flowering time of the indicated genotypes and isogenic lines derived from the introgression of *SUC2::HA:CO* transgenes into *aba1-6* or *gi* mutant backgrounds. Note that for the *SUC2::HA:CO aba1-6 gi-100* combination, two sibling lines were analysed. An ANOVA test to assess the impact of mutations at the *ABA1* and *GI* loci on leaf number at bolting was run separately for transgenic (*SUC2::HA:CO*) and control plants (vertical bar). For both groups, genotype was a statistically significant predictor of leaf number at bolting (*p* < 0.001). Letters at the top of boxplots indicate if genotypes showed statistically significant differences (*p* < 0.05) according to a Tukey post-hoc test. The number of samples analysed for each genotype is shown at the bottom of the graph. (B) GI ChiPseq peaks on the *FT* region coincide with DNAse accessible chromatin (referred to as enhancer blocks C and E). CO recruitment to the CORE region is regulated by ABA and GI (thick arrows). ABA and GI might act independently to promote CO binding to the CORE region. ABA might also influence the accessibility of GI to Block C/E (dotted arrows), thus indirectly controlling CO recruitment.

## DISCUSSION

Our study describes a yet uncharacterised role for ABA on CO function. Since ABA biosynthesis and signalling is enriched in the phloem companion cells of the leaf (Mustilli *et al*., 2002; Kuromori *et al*., 2014), endogenous CO function might be subject to varying ABA accumulation, which is usually related to water deficit status. Our experiments do not imply a direct role of ABA in controlling CO function as this could be mediated by one or more CO-interacting protein(s) or factors that are not present in ABA-deficient plants. However, they establish a role for ABA and its signalling components in influencing *FT* promoter accessibility to CO, particularly the *CORE1/2* region, subsequently leading to RNAPol II engagement at the *FT 5’ UTR*.

Recent evidence supports the role of GI as a potential target of ABA action during DE. First, we previously highlighted the key role of GI in enhancing *FT* expression in response to ABA, and that GI function was sensitive to the ABA signalling status (Riboni *et al*., 2016). Second, the epistatic interaction between *gi* and *aba1* mutants in the regulation of flowering time (bolting time in particular) of *SUC2::HA:CO* plants aligns with a model where ABA and GI could act in the same pathway to regulate CO recruitment to the *FT* promoter, although the regulatory role of ABA on GI function remains unknown. GI was shown to stimulate ABA production which might suggest an indirect effect on CO recruitment exerted by GI (Baek *et al*., 2020). However, we reported an increased levels of ABA sensitivity in *gi* plants under controlled water deficit conditions (Siemiatkowska *et al*., 2022), suggesting that GI might be crucial for relaying ABA signalling upstream of *FT*. GI exerts a direct influence on *FT* activation by associating to its promoter (Sawa and Kay, 2011). To further define the sites of GI binding onto the *FT* regulatory region, we analysed GI ChIPseq data describing the genome-wide occupancy of GI (Nohales *et al*., 2019). GI binding regions on the *FT* promoter (Fig. 6B) closely overlap with two DNAse accessible chromatin locations (Zhang *et al*., 2012). These regions were functionally defined as block C and E enhancers (Adrian *et al*., 2010; Liu *et al*., 2014; Zicola *et al*., 2019). Block C contains NF-Y binding sites, which facilitate looping with the *CORE* region and consequent *FT* transcriptional activation (Cao *et al*., 2014; Siriwardana *et al*., 2016). Block E sequence exhibits binding sites for different transcriptional regulators, including the *FT* repressor SCHLAFMUTZE (SMZ), which has been shown to bind to GI (Mathieu *et al*., 2009; Sawa and Kay, 2011). While GI is unlikely to bind DNA and despite its molecular function is still unclear, it is required to control transcriptional events through direct interaction with specific transcriptional regulators (Sawa and Kay, 2011; Song *et al*., 2014; Kubota *et al*., 2017), possibly by acting as a general chaperone-like protein (Cha *et al*., 2017). These observations could point to a model whereby GI controls CO recruitment through binding to Block E and C enhancers, perhaps by affecting local chromatin conformation or CO protein complex accessibility (Fig.6B).

Different NF-Y encoding genes act as regulators of ABA transcriptional responses (Kumimoto *et al*., 2013; Song *et al*., 2016) and *nf-yb2* mutants display an exaggerated delay in flowering time compared to wild-type controls under osmotic stress conditions (Chen *et al*., 2007). Considering the mild flowering time defects of *nf-yb2* mutants under normal growth conditions (Kumimoto *et al*., 2008), this observation may point to a specific requirement for *NF-YB2* subunit in promoting flowering under osmotic stress, which is ABA regulated. A similar observation about the requirement for NF-Y in mediating ABA-dependent flowering derives from the lack of DE of *nf-yc3 yc4 yc9* mutants (Hwang *et al*., 2019). Because NF-YB/C can form complexes with both CO and NF-YA subunits, their contribution in relaying ABA signals may depend on either CO, NF-YA or both functions. Moreover, NF-YC subunits interact with the ABFs bZIP to promote the transcriptional activation of *SOC1* (Hwang *et al*., 2019). Whether this model of interplay between ABFs and NF-YA/B/C can be applied to the *FT* promoter regulation is unclear mainly because the ABFs do not appear to bind to the *FT* chromatin region (Song *et al*., 2016; Hwang *et al*., 2019). Intriguingly, recent data support the physical interaction between CO and ABF1/2/3/4 and their antagonistic roles at salinity-responsive genes promoters (Du *et al*., 2023). Our ChIPseq analyses revealed an over-representation of genes related to biotic and abiotic responses (particularly salt and water stress response) among *bona fide* CO direct targets. This points to a broader role for CO in influencing processes unrelated to flowering time regulation in Arabidopsis and other species (Du *et al*., 2023; Biancucci *et al*., 2024, Preprint). It might also suggest a potential positive feedback mechanism, in which ABA mediates between abiotic stress levels and CO occupancy at regulated promoters. Previous reports have described direct links between ABA signalling and function of chromatin regulatory complexes that affect *FT* expression (Farrona *et al*., 2004; Peirats-Llobet *et al*., 2016; Katagiri *et al*., 2024). Here, genetic, and molecular evidence suggests that ABA might also act through GI-mediated recruitment of CO to regulate *FT* expression. Therefore, our data and experimental approaches offer new tools for investigating the ABA-dependent level of regulation of photoperiodic flowering.

## Acknowledgments

We thank personnel at Orto Botanico Città Studi and Viktoriia Kyrychenko for plant care, Fabio Fornara and Federico Zambelli (University of Milan) for insightful comments and guidance on ChIPseq analysis, respectively. This study has been supported by a grant from the Italian Ministerial of Research PRIN 2022 “Light and drought signals integration driving development transitions and adaptations in plants – LIDS” Ref: 2022T2737Y and by a research grant from the HFSP Ref: RGP0011/2019. DM is supported by a research fellowship co-funded by the European Union – ESF, REACT-EU, PON Ricerca e Innovazione 2014-2020. The funders had no role in study design, data collection and analysis, decision to publish, or preparation of the manuscript.

## Author contribution

ART, GP, LC: conceptualization and investigation; AS, SC, PKK, GC, TX, EV, DM: investigation; BL: formal analysis; GP, ART, LC: writing – original draft; EK, MG, CT, LC: writing – review & editing; LC: supervision; LC: funding acquisition

**No conflict of interest declared**

