## Supplementary material for "Abscisic acid and GIGANTEA signalling converge to regulate the recruitment of CONSTANS to the *FT* promoter and activate floral transition": Suppl File Figures

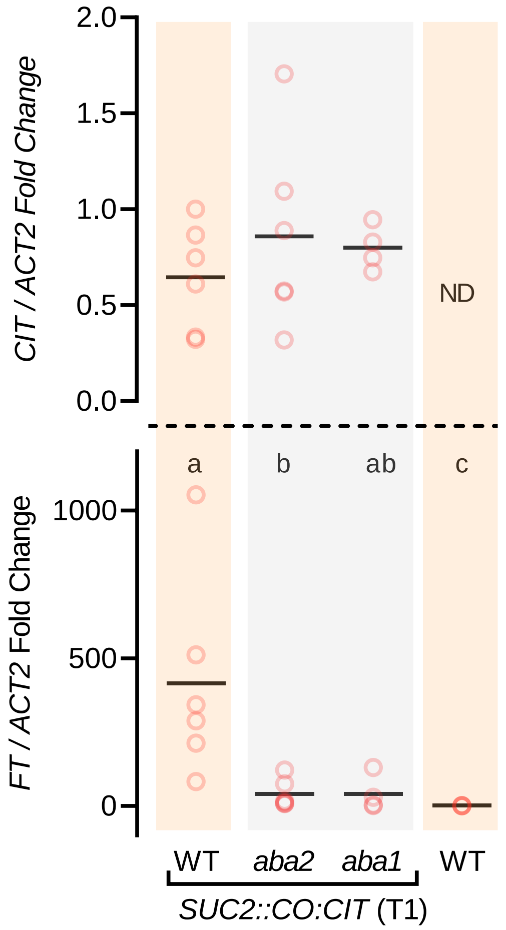


**Supplementary Figure S1. ABA promotes CO function and *FT* expression.**

Real-Time qPCR detection of *CITRINE (CIT,* Top panel) and *FT* (bottom panel) transcripts accumulation. Horizontal Lines represent the mean fold change of *ACT2*-normalised *CIT*/*FT* accumulation. Each dot corresponds to a single 20-day-old T1 *SUC2::CO:CIT* transgenic plant (whole rosette, n = 4 - 7) or fully expanded leaves of the wild type (empty vector control, n = 3) harvested at ZT12. Values are relative to the geometric mean of the wild type (*FT* expression) or *SUC2::CO:CIT* (*CO:CIT* expression). ND = not determined. Plants were grown on soil under LDs. Letters at the top indicate statistically significant differences ($p$ < 0.05) according to a Tukey post-hoc test.

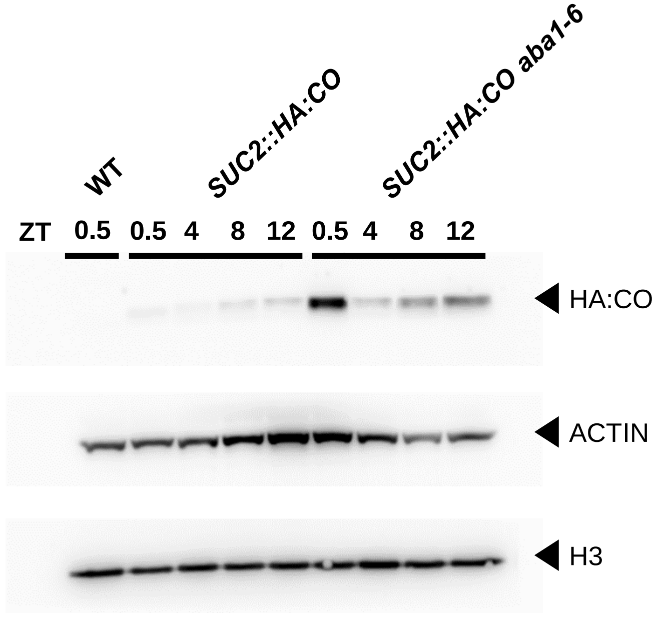


**Supplementary Figure S2. CO accumulation in ABA deficient mutants during the light phase.**

Immunoblot detection of HA:CO protein from total lysates derived from *SUC2::HA:CO* plants (WT or *aba1-6* backgrounds) assayed at the indicated ZT points under LDs. WT, non-transgenic plants served as a negative control, whereas the detection of ACTIN and Histone 3 (H3) afforded a loading controls.

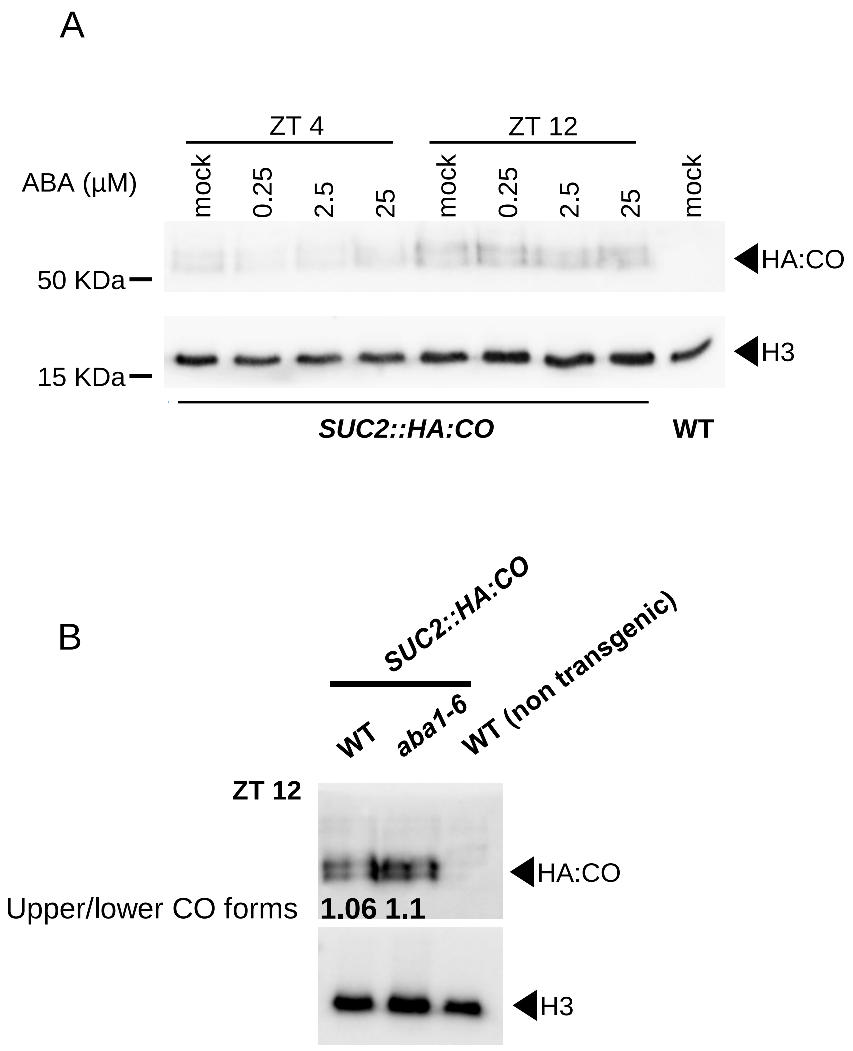


**Supplementary Figure S3. ABA does not alter CO accumulation or modifications.**

(A) Immunoblot detection of HA:CO protein from nuclear fractions derived from *SUC2::HA:CO* plants assayed at ZT4 and 12 under LDs. WT, non-transgenic plants served as a negative control, whereas the detection of Histone 3 (H3) afforded a loading control. Numbers refer to molecular mass according to the migration of a protein ladder. ABA (or mock solution carrying an appropriate amount of EtOH, the diluent for ABA) was applied at different concentrations (µM) on soil (B) Immunoblot detection of HA:CO protein from nuclear fractions derived from *SUC2::HA:CO* (WT or *aba1-6*) plants assayed at ZT12 under LDs. Numbers below HA:CO band represent the ratio of the intensity of the upper and lower bands.

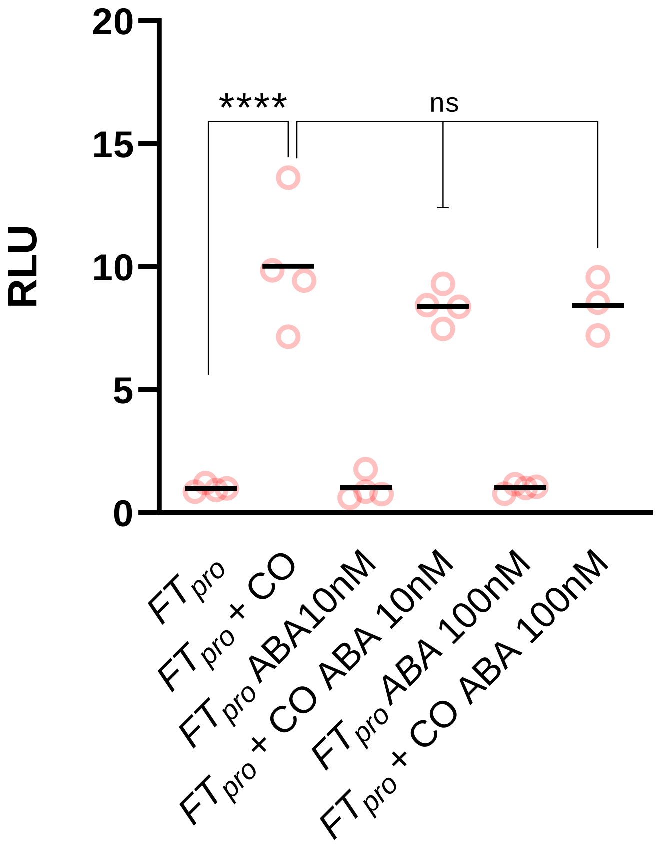


**Supplementary Figure S4. ABA does not affect CO-dependent activation of the *FT* promoter in protoplasts transient transformation.** Relative Light Units (RLU) for Luc/Ren activities measured in protoplasts transformed with combinations of *FTpro::LUC* reporter, *35S::CO* after incubations with different ABA concentrations. Asterisks denote a significant difference ($p < 0.001$), ns = not significant

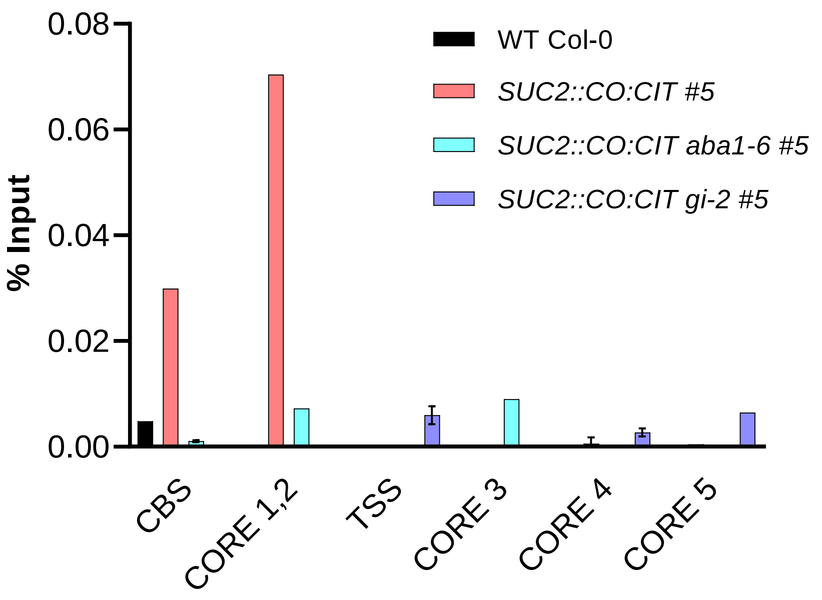


**Supplementary Figure S5. Biological replication of CO enrichment on the *FT* promoter.**

Relative enrichment of CO:CIT on the *FT* promoter in isogenic lines *SUC2::CO:CIT#5* (WT *aba1-6* and *gi-2* mutant backgrounds). Non transgenic, WT (Col-0) was used as negative control. Bars are the mean ± SE (n = 4 technical replicates).

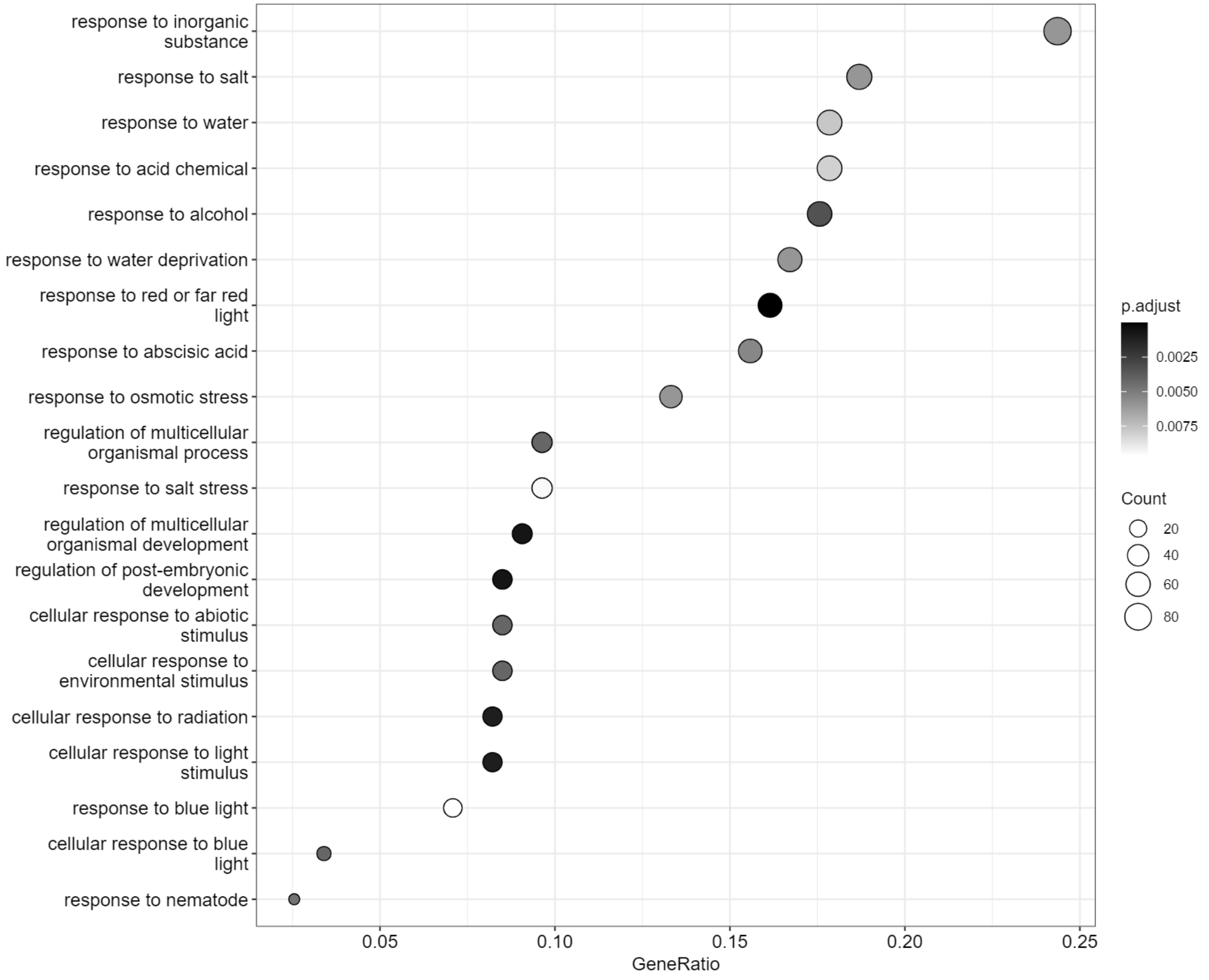


**Figure S6. Gene-set enrichment plot for biological process terms**

Data represent the functional classification of the Intersection between genes falling next to *35S::CO* peaks and upregulated DEGs in *35S::CO* (data from de los Reyes et al., 2024).

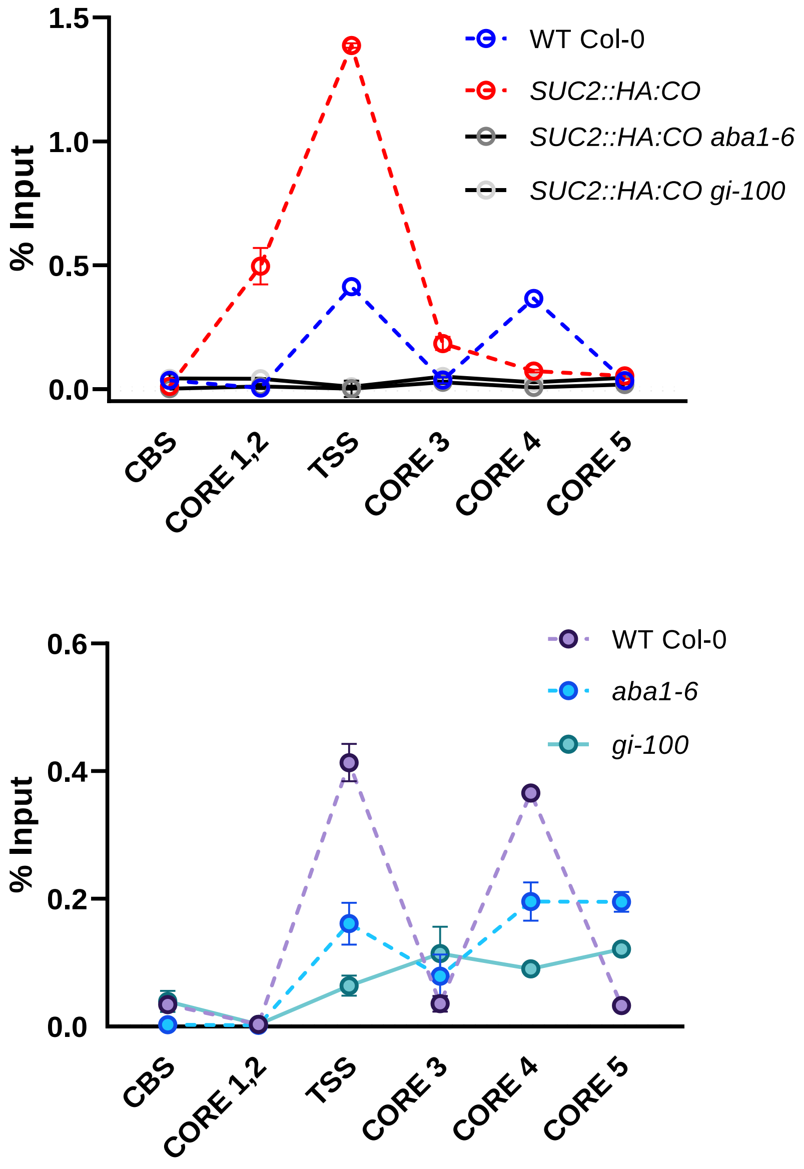


**Supplementary Figure S7. Biological replication of RNAPol II enrichment on the *FT* promoter.**

RNAPolII deposition (expressed as % of input) on the *FT* locus in the indicated genotypes. The same experiment was plotted in two separate graphs to emphasize the effect of *SUC2::HA:CO* on RNAPol II recruitment. Each point represents the mean ± SE (n = 4 technical replicates).

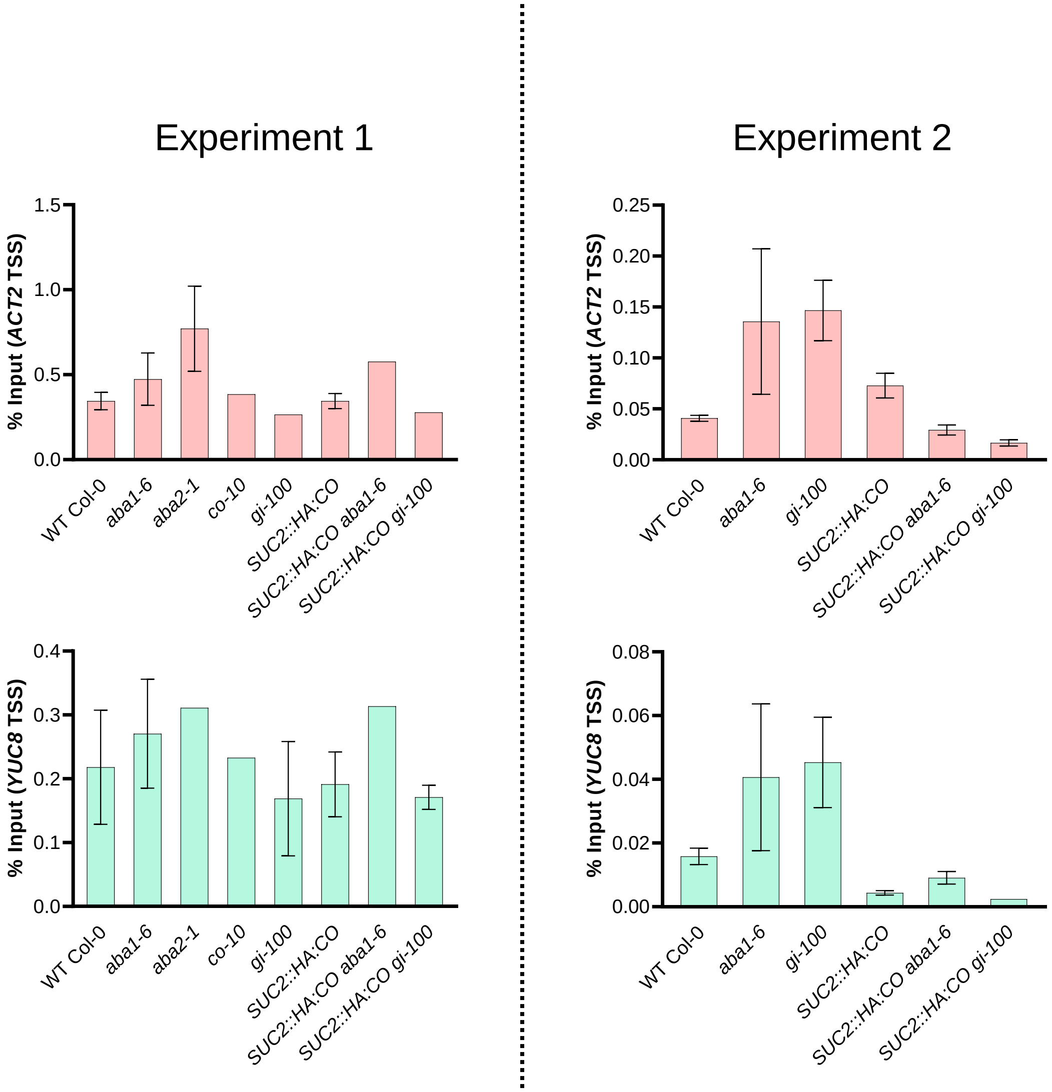


**Supplementary Figure S8. RNAPol II recruitment to the *TSS* of on *YUC8* and *ACT2* is not affected in ABA deficient or *gi* backgrounds.**

RNAPol II deposition on the *TSS* region (expressed as % of input) of the *ACT2 and YUC8* loci in the indicated genotypes (experiments). Histograms represent the mean ± SE (n = 4 technical replicates).

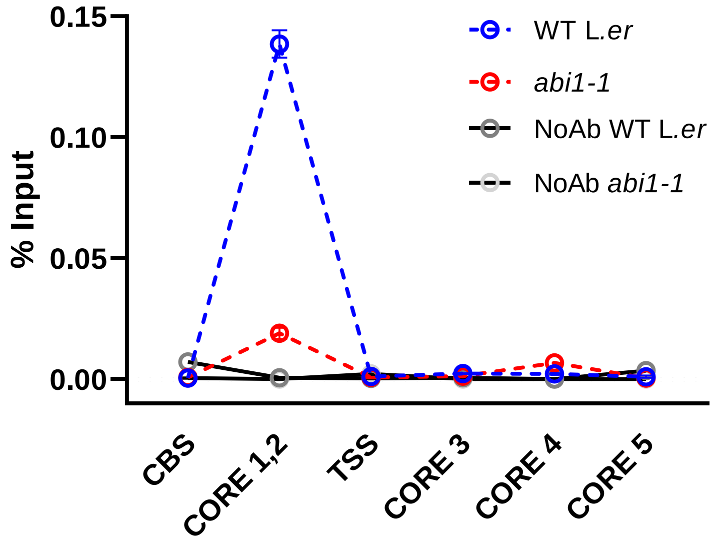


**Supplementary Figure S9. Biological replication of RNAPol II enrichment on the *FT* promoter.**

RNAPol II deposition (expressed as % of input) on the *FT* locus in the WT (L.*er*) and *abi1-1* mutant. No antibody denotes a negative control for the ChIP assay. Each point represents the mean ± SE (n = 4 technical replicates).

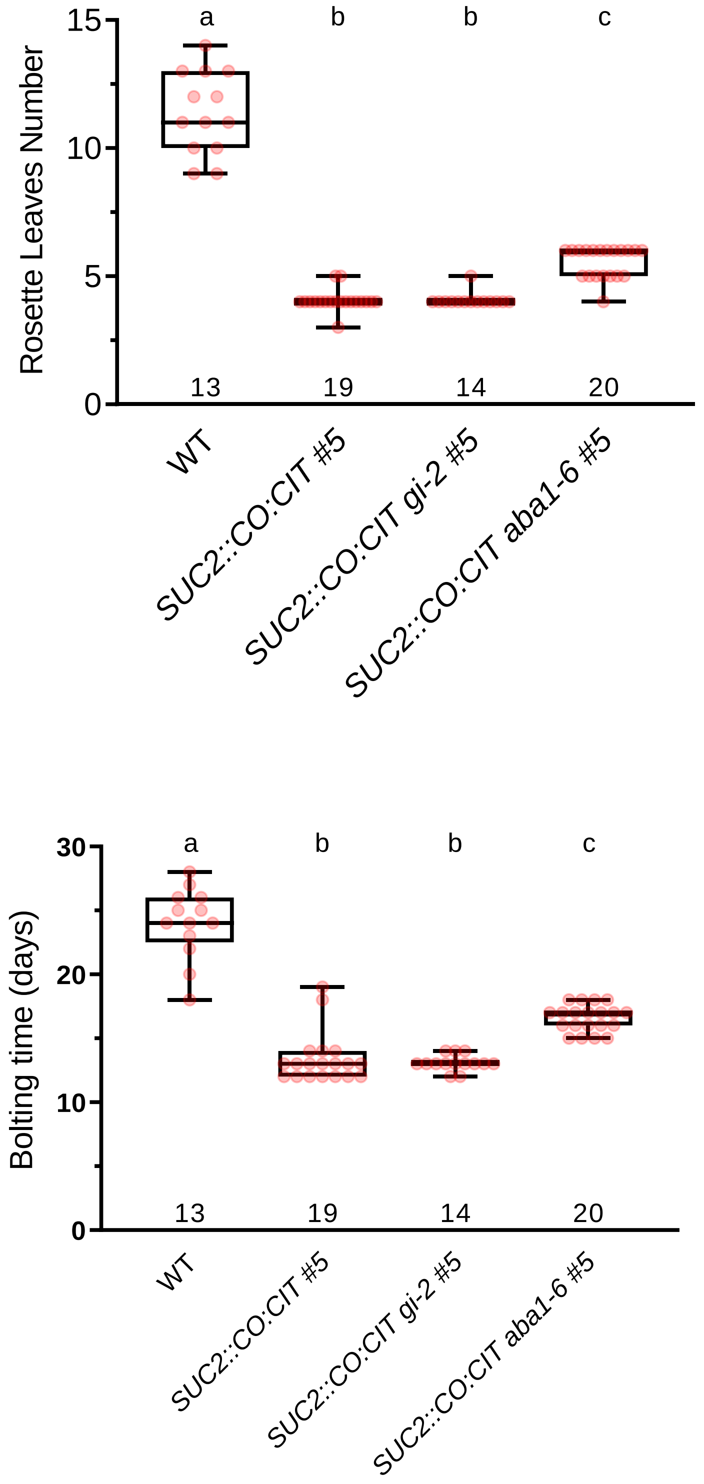


**Supplementary Figure S10. *gi* is insufficient to suppress *SUC2::CO:CIT*.** Boxplot of flowering time (rosette leaves number, top panel and bolting time, bottom panel) for the indicated genotypes. Isogenic lines *SUC2::CO:CIT #5 aba1-6* were included as a control. Different letters denote different ($p < 0.05$) groups according to a Tukey post-hoc tests. The number of samples analysed for each genotype is shown at the bottom of the graph.

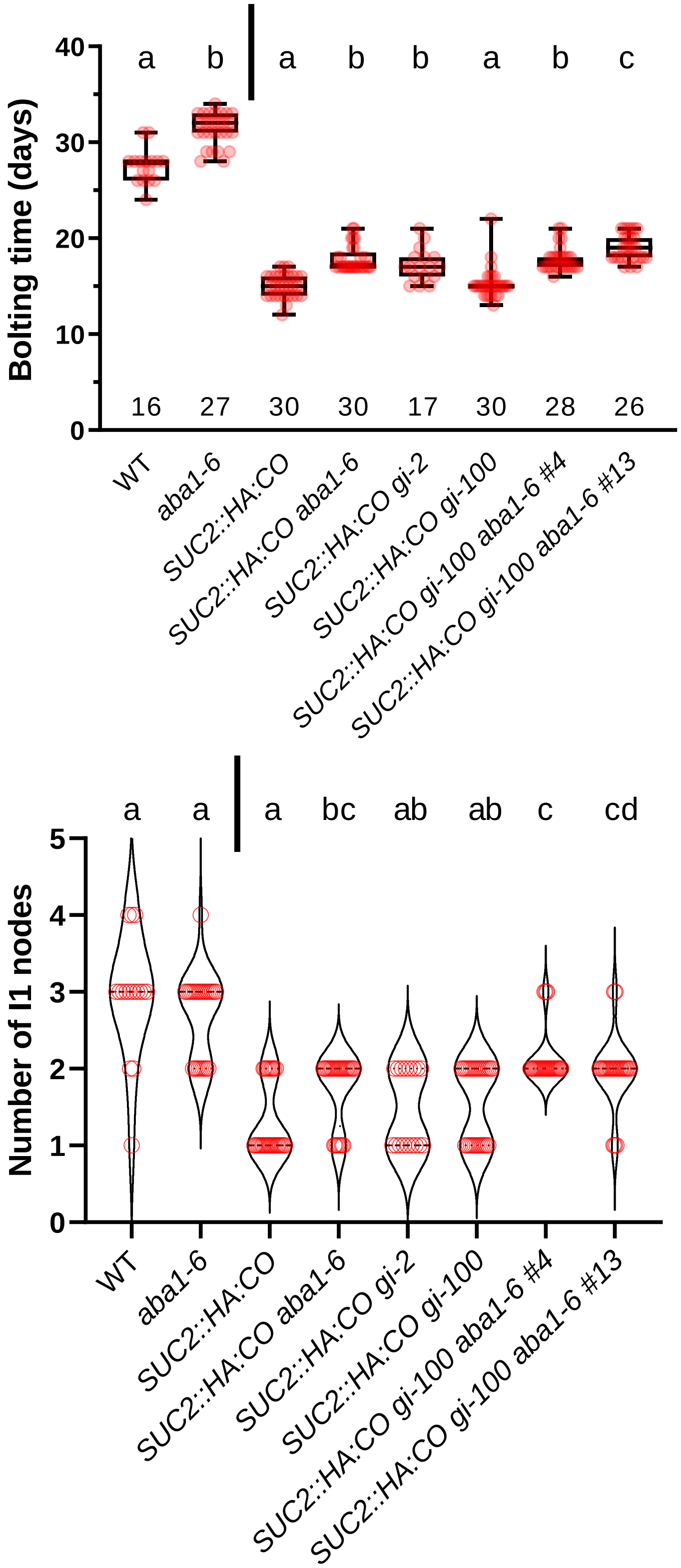


**Supplementary Figure S11. Effects of *gi* and *aba1* mutations in attenuating of *SUC2::HA:CO*.** Boxplot of bolting time (top) and violin plot of number of I1 nodes (bottom) of the indicated genotypes and isogenic lines derived from the introgression of *SUC2::HA:CO* transgenes into *aba1-6* and/or *gi* mutant backgrounds. ANOVA tests to assess the impact of mutations at the ABA and GI loci on days to bolting and number of I1 nodes were run separately for transgenic (*SUC2::HA:CO*) and non-transgenic plants (vertical bar). Letters at the top of boxplots indicate if genotypes showed statistically significant differences ($p < 0.05$) according to a Tukey post-hoc test. The number of samples analysed for each genotype is shown at the bottom of the graph.
